# Intracellular Ebola Virus nucleocapsid assembly revealed by *in situ* cryo-electron tomography

**DOI:** 10.1101/2023.12.01.569666

**Authors:** Reika Watanabe, Dawid Zyla, Diptiben Parekh, Connor Hong, Ying Jones, Sharon L. Schendel, Willian Wan, Guillaume Castillon, Erica Ollmann Saphire

## Abstract

Filoviruses, including Ebola and Marburg viruses, cause hemorrhagic fevers with up to 90% lethality. The viral nucleocapsid is assembled by polymerization of the nucleoprotein (NP) along the viral genome, together with the viral proteins VP24 and VP35. We employed cryo-electron tomography of cells transfected with viral proteins and infected with model Ebola virus, to illuminate assembly intermediates as well as a 9Å map of the complete intracellular assembly. This structure reveals a previously unresolved, third, and outer layer of NP complexed with VP35. The intrinsically-disordered-region together with the C-terminal domain of this outer layer of NP provides the constant-width between intracellular nucleocapsid bundles and likely functions as a flexible tether to the viral matrix protein in virion. A comparison of intracellular nucleocapsid with prior in-virion nucleocapsid structures reveals the nucleocapsid further condenses vertically in-virion. The interfaces responsible for nucleocapsid assembly are highly conserved and offer targets for broadly effective antivirals.

## Introduction

Filoviruses, including several species of Ebola and Marburg viruses, cause severe hemorrhagic fever in humans^1^. A vaccine is available for just one of these viral species^2,3^. Antibody treatments are also approved, but also, thus far for just one species. Further, these viruses can lurk in immune-privileged sites, and emerge 1-2 years later to trigger new chains of infection^4^. Development of antiviral drugs active against a range of filoviruses and which can access immune-privileged sites is a high priority. To meet this objective, we must understand the critical and conserved structures that are indispensable for viral assembly and replication.

Filoviruses belong to the order *Mononegavirales*, along with other severe human pathogens such as rabies, measles, and respiratory syncytial viruses. *Mononegavirales* are enveloped viruses with a non-segmented, single-stranded, negative-sense RNA genome that is encapsulated by nucleoprotein (NP) to form a helical nucleocapsid. In filoviruses, two other viral proteins, VP35 and VP24, are also required to form the nucleocapsid^5,6^. Binding of VP35 to nascent NP is thought to prevent oligomerization of NP before binding to the viral RNA genome^7–12^. VP35 also functions as an essential virus RNA polymerase cofactor, linking the viral polymerase with the genome-NP template^13–15^. VP24 is unique to the filovirus family. In addition to its critical role in nucleocapsid formation, VP24 regulates viral transcription and replication^16^ and intracellular transport of the nucleocapsid^17^. Importantly, VP35 and VP24 also suppress host immune signaling, influencing viral replication and pathogenicity^18–23^. Prior work revealed nucleocapsid-like structures in budded viruses or virus-like-particles^24,25^. In these structures, a repeating unit of the nucleocapsid consists of two adjacent copies of core NP molecules that polymerizes along the viral genome, plus two VP24 molecules^25^. The N-terminal half of NP was visible, but the location of both the C-terminal half of NP and the VP35 protein remained unknown.

Until recently, it was impossible to reveal protein structures directly inside of virus-infected cells, except very thin peripheral parts of cells or budded virus particles. Therefore, we had no structural insights into how the Ebola virus replicates and how nucleocapsids assemble in infected cells prior to budding and release. Here, we use cryo-focused ion beam (FIB)-enabled cryo-electron tomography (ET) to directly visualize Ebola virus nucleocapsid assembly inside cells. We used both cells transfected with viral proteins (allowing analysis of individual components as well as truncation mutants) as well as cells in the context of viral infection using a biologically contained Ebola virus model. We identify distinct intermediate nucleocapsid structures assembled in the host cytoplasm and provide the first molecular model of a fully assembled Ebola nucleocapsid prior to viral egress. Importantly, a comparison of this intracellular structure to prior post-egress structures reveals conformational changes in the nucleocapsid that occur upon virion incorporation. Further, the analysis here also revealed an additional outer layer of NP in the nucleocapsid, held RNA-free by VP35 and which likely is what mediates the flexible tether to the viral membrane. The molecular interaction sites that build these assemblies are conserved among filoviruses and provide the templates for design of broadly effective antivirals.

## Results

### Identification of nucleocapsid assembly intermediates in cells expressing NP, VP24, and VP35

To elucidate the intracellular nucleocapsid assembly process, we first used HEK 293T cells transfected with complete EBOV NP or NP lacking C-terminal amino acids 601-739 (NPΔ601-739), in combination with VP24 and VP35. NPΔ601-739 was selected for analysis as it is the shortest NP known to form a fully assembled nucleocapsid^26^. We expected that comparing two nucleocapsid structures with full-length or C-terminally truncated NP would help identify the previously unassigned C-terminal domain of NP^25^ in the intracellular nucleocapsid structure. Immunofluorescence microscopy confirms that all transfected proteins (NP, VP35, and VP24) co-localized in cytoplasm (Figures S1A-S1F). Resin-embedded, ultra-thin section transmission electron microscopy shows that in the presence of VP24 and VP35, both NP and NPΔ601-739 efficiently form bundles of electron-opaque, assembled, intracellular nucleocapsid-like structures inside the cytoplasm in a region that excludes the majority of subcellular organelles, similar to EBOV-GFP-ΔVP30-infected cells (Figures S1G-S1P).

To gain structural insight into the nucleocapsid assembly process, we investigated the transfected cells using cryo-electron tomography coupled with focused ion beam milling^27^. In cytoplasm of full-length NP-, VP24- and VP35-expressing cells, we identified three distinct intracellular nucleocapsid precursors (Figure 1): (1) loosely coiled NP polymers, which resemble purified full-length EBOV NP bound to cellular RNA from mammalian cells^24^, (2) regionally condensed NP polymers, whose condensed parts resemble purified C-terminally truncated EBOV NP (amino acids 1-451) bound to cellular RNA from mammalian cells^24,28^ and (3) fully assembled nucleocapsid-like structures, which resemble nucleocapsids found in Ebola virions and virus-like particles^24,25^ (Figure 1).

**Figure 1.**
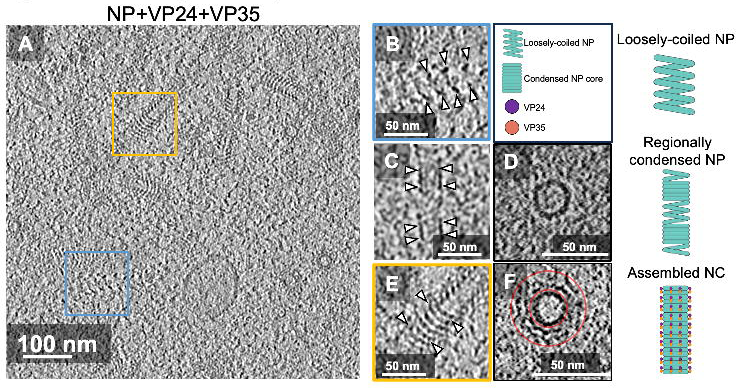
Ebola nucleocapsid assembly intermediates. (A-F) Representative tomographic slice of cells transiently expressing NP, VP24, and VP35 (A); enlarged views showing loosely-coiled NP (B); regionally condensed NP-only coils (C, D), and fully assembled nucleocapsid with additional outer layers on condensed NP (E, F), highlighted by the arrowheads respectively. Schematic representations of the intermediate structures are shown on the right. Cross-sectional views of (C and E) are shown in (D and F), respectively. The cross-sectional diameter of NP (D) is 23 nm, and the assembled nucleocapsid (F) is ∼50 nm. The outer layer of nucleocapsid lies between the red circles in the cross-sectional view (F). See also Figures S1, and Video S1.

### Structure of intracellular Ebola NP**Δ**601-739-VP24-VP35 nucleocapsid

We determined fully assembled nucleocapsid coordinates by cylinder correlation, and performed subtomogram averaging to resolve overall assembled nucleocapsid structures (Figures S2, and Videos S1-S2). We found that both intracellular nucleocapsid-like structures (whether from complete NP or C-terminally truncated NP) form a left-handed helix with a diameter of 52 nm and nearly identical helical parameters (Figures 2B, 3A). To obtain higher resolution structures of individual repeating units that form these helical structures, we performed further subtomogram analysis by re-extracting subtomograms, and centering their repeating units based on the helical parameters (Figure S2). We performed 3D classification and 3D refinement without applying any helical symmetry, considering the highly heterogeneous nature of filovirus nucleocapsid structures in virions^24,25,29^. We obtained 9 Å and 18 Å resolution maps of intracellular nucleocapsid structures in cells expressing C-terminally truncated NPΔ601-739 and full-length NP, respectively (Figures 2C-2E, 2L, 3B-3C, S2). Our 9 Å map shows its highest local resolution in the central repeating unit (Figure 2L). By combining this map with pre-existing high-resolution structures and flexible fitting, we could build a molecular model of the intracellular nucleocapsid structure (Figures 2F-2I, S3, summarized in Video S3).

**Figure 2.**
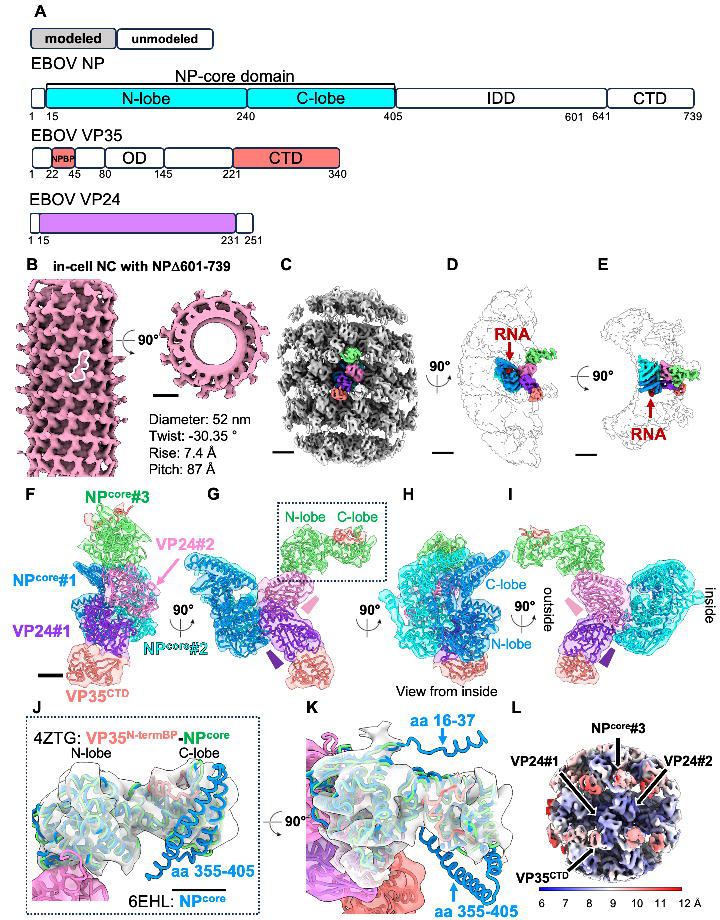
Structure of intracellular Ebola NPΔ601-739-VP24-VP35 nucleocapsid. (A) Domain organization of Ebola virus (EBOV) NP, VP35, and VP24. Regions that have been modeled and unmodeled in this study are indicated by filled and open boxes, respectively. (B) The initial map of intracellular Ebola NPΔ601-739-VP24-VP35 nucleocapsid (pink). The repeating unit is marked with a white outline. (C-E) Intracellular NPΔ601-739-VP24-VP35 nucleocapsid subunit densities. The two central NP core domains (NP^core^#1, NP^core^#2) are in blue and cyan. Two copies of VP24 (VP24#1, VP24#2) are in purple and pink. The C-terminal domain of VP35 (VP35^CTD^) is in orange, and the third NP core (NP^core^#3) is in green. Continuous red density at the interior core of the NP copies corresponds to RNA. (F-I) Molecular models fitted into EM densities. Colors are as in (C-E). Pink and purple cones indicate the orientation of the pyramid shape of VP24 molecules^18^ in (G, I). The locations of N-lobe (aa 15-240) and C-lobe (aa 241-405) of NP are shown in (G, J). (J and K) Crystal structure of NP complexed with the VP35 N-terminal NP-binding peptide (green and orange; PDB: 4ZTG)^11^ and the NP core structure obtained from isolated NP (aa 16-405)-RNA helix (blue; PDB: 6EHL)^25^ each fitted into the outermost density indicated by the dotted line in (G). (L) Local resolution map. Scale bars indicate 10 nm (B), 5 nm (C-E), 2 nm (F, J). See also Figures S2, S3 and Videos S2-S3.

Within a single repeating unit (colored in Figures 2C-2E), we assigned two NP core domains from PDB: 6EHL^25^ (NP^core^#1 in blue, NP^core^#2 in cyan) and two VP24 molecules from PDB: 4M0Q^30^ (VP24 #1 in purple, VP24 #2 in pink)(Figures 2F-2I). Pink and purple cones in (G and I) indicate the orientation of the pyramid shape of VP24 molecules^18^. This model of a pair of RNA-bound NPs and their antiparallel VP24s (one inward and one outerward) match well with the previous in-virion nucleocapsid model obtained with Ebola virus viral-like-particles and chemically fixed Ebola virions^25^. In addition, we could now also visualize an outer layer of density of the intracellular nucleocapsid map and assign this density to a repeating third copy of NP (Figures 2F-2I). This newly-assigned NP (NP^core^#3 in green) is monomeric in nature (not attached to other copies of NP) and is attached to VP24#2.

Serial docking of this newly assigned outer NP density with previous in-virion cryo-EM and X-ray structures [NP core (PDB: 6EHL)^25^ and NP bound to VP35 N-terminal NP-binding peptide (PDB: 4ZTG^11^, 4YPI^12^)], showed the best in-map density correlation with a crystal structure of NP (aa 36-354 in green) in complex with the N-terminal NP-binding peptide of VP35 (aa 22-45 in orange) (PDB: 4ZTG, in green and orange)^11^ (Figures 2J, 2K). This NP-binding peptide of VP35 holds NP in a monomeric state^11,12^. Indeed, the N-terminus NP binding peptide of VP35 (aa 22-45 in orange) is recognizable and attached to the outer copy of NP in our cryo-ET map (Figures 2J, 2K, S3F, S3G and Video S3).

In contrast, attempted fitting of the conformation of NP that forms the central nucleocapsid coil (PDB: 6EHL, in blue) into the same outer density is poor (Figures 2J, and 2K in blue). The N-terminal alpha-helix (aa 16-37), part of the C-terminal penultimate helix, and the entire long clamp helix (aa 355-405) are well outside the cryo-ET density for NP#3 (Figures 2J, and 2K in blue). These three helices are disordered in monomeric VP35-bound NP^11^ and are only stabilized when NPs instead bind to RNA and polymerize into helices^25^.

Taken together, our EM map reveals not only the presence of an unexpected third NP molecule in each repeating unit, but also that two different states of NP molecules simultaneously co-exist in fully assembled nucleocapsids: one as an oligomeric helical form bound to RNA, and the other held RNA-free and monomeric by VP35 NP-binding peptide. Interestingly, superimposition of one of the central NP-VP24 subunits, NP#1-VP24#1 with this newly annotated VP24#2-outer NP#3 shows an almost perfect match of both VP24 and NP N-lobes, demonstrating that newly identified NP#3 binds to VP24#2 using the same interface as NP#1-VP24#1 (Figure 6F, and Video S3). In previous cryo-ET on budded virions, it was clear that adjacent VP24s #1 and #2 were oriented in opposite directions, VP24#1 facing “in” and VP24#2 facing “out”, but it was not yet clear why they would be organized in this way^25^ (Figures 2F-2I). In virion, there is additional density in contact with the outward-facing VP24 that was not previously assignable. In light of the new in-cell model, we propose that this density belongs to the third copy of NP.

On the outer surface of the nucleocapsid, in addition to the short N-terminal NP-binding peptide of VP35, we can also predict the position of the C-terminal domain of VP35 by fitting its crystal structure (VP35^CTD^, amino acids 221-340, PDB: 3FKE^31^; Figures 2F-2I, S3H-S3J, and Video S3) in the other smaller outer densities, associating with VP24#1.

### C-terminal regions of NP are located between intracellular nucleocapsids

Our two in-cell nucleocapsid-like structures, one for C-terminally truncated NPΔ601-739 and the other for full-length NP, are essentially identical, except in resolution (9 Å vs. 18 Å; Figures 2C-2E vs 3B-3C and S4A-S4L). The molecular model for C-terminally truncated NP fits well into the 18 Å map of full-length NP (Figures 3D-3E) with no unoccupied density, suggesting that the C-terminal domain of NP is not visible in the current 18 Å map of the intracellular nucleocapsid formed by full-length NP. In the cytoplasm, intracellular nucleocapsids from both full-length and C-terminally truncated NPΔ601-739 often form parallel bundles (Figures 3G, 3H, 3J, 3K and Videos S1 and S2) similarly to authentic Ebola virus infected cells^5,32,33^. We measured the center-to-center distances of nucleocapsids in these bundles based on the coordinates obtained from filament tracing (Figures S2). Both nucleocapsids have an identical 52 nm diameter, but the bundles formed by full-length NP exhibit greater nucleocapsid-to-nucleocapsid distance (14 nm, 66-52=14; Figure 3I) than do bundles formed by NPΔ601-739 (2 nm, 54-52=2; Figure 3L). By combining the location of the C-terminal end of NP^core^#3 on the outer layer with the difference in nucleocapsid-to-nucleocapsid distance, we propose that the C-terminal domain of NP (NP^CTD^, amino acids 641-739), and its intrinsically disordered linker to the core (aa 601-644) occupies the gap between the nucleocapsids in the bundles (Figure 3F). Moreover the nucleocapsid is always centered in the gap, with a consistent 7 nm spacing on either side of the 52 nm-wide nucleocapsid (7+7=14 nm; Figures 3F-3I, and 7B).

**Figure 3.**
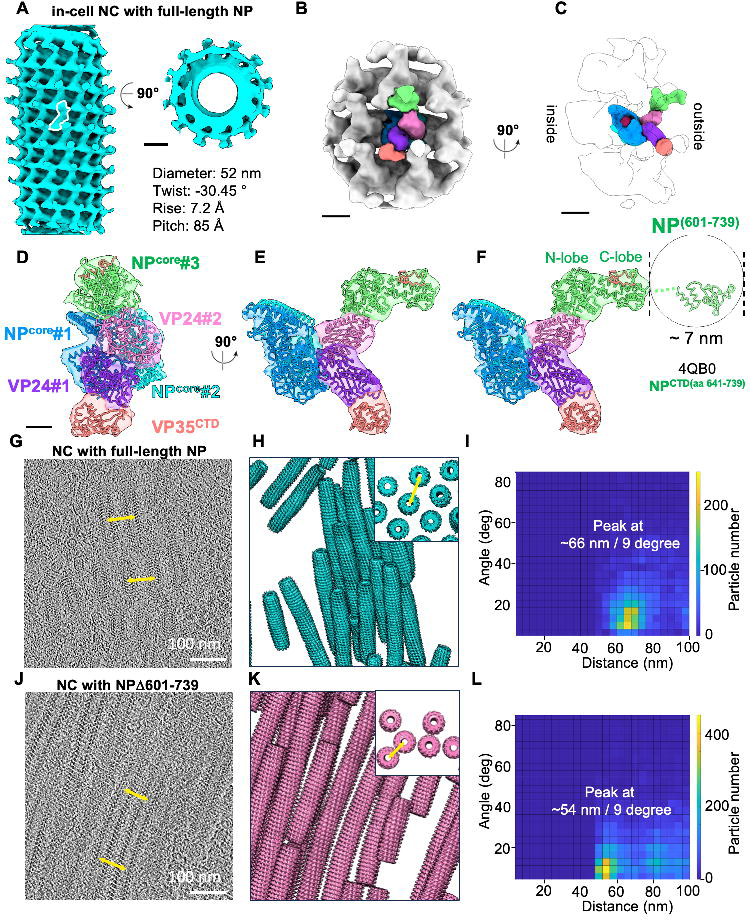
Structure of intracellular Ebola NP-VP24-VP35 from full-length NP. (A) Initial map of the intracellular Ebola virus nucleocapsid made from full-length NP-VP24-VP35. The repeating unit is marked with a white outline. (B-C), Intracellular full-length NP-VP24-VP35 nucleocapsid subunit densities. Density colors correspond to those for Figure 2. (D-F) Molecular models fitted into the EM densities and the hypothetical position of NP aa 601-739 based on the measured distance between intracellular nucleocapsids. The crystal structure of the NP C-terminal domain (aa 641-739; PDB: 4QB0)^58^ is fit into the gap. (G and J) Representative tomographic slices of cells, (H and K) Initial averages mapped back onto tomograms, and (I and L) Two-dimensional histograms showing the distance of the center-to-center and angle of the intracellular nucleocapsids formed by full-length NP-VP24-VP35 and NPΔ601-739-VP24-VP35, respectively. Yellow lines in G, H, J and K indicate the center-to-center distance of the closest neighboring nucleocapsid. H and K panel insets show oriented views to visualize the distance between aligned nucleocapsids. Scale bars indicate 10 nm (A), 5 nm (B-C), 2 nm (D), 100 nm (G, J). See also Figures S2 and Video S1-S2.

### Molecular model of in-virion nucleocapsids and condensation of intracellular nucleocapsid upon virion incorporation

In previous studies of in-virion Ebola virus nucleocapsids, two distal outer densities were present but could not be assigned^25^. Docking of our in-cell nucleocapsid molecular model into a map generated by previous EM analysis of nucleocapsid in virus-like particles (VLPs) (EMD-3871)^25^, followed by flexible refinement, yielded a complete *in-virion* nucleocapsid model for Ebola virus (Figures 4A-4C). This updated in-virion nucleocapsid model also fits well into a previous EM map of the nucleocapsid in a chemically fixed, authentic Ebola virus (EMD-3873)^25^ (Figures 4D-4F). In authentic virions, the density for the VP35^CTD^ is very weak, suggesting that this position is rarely fully or stably occupied in the virus^25^.

**Figure 4.**
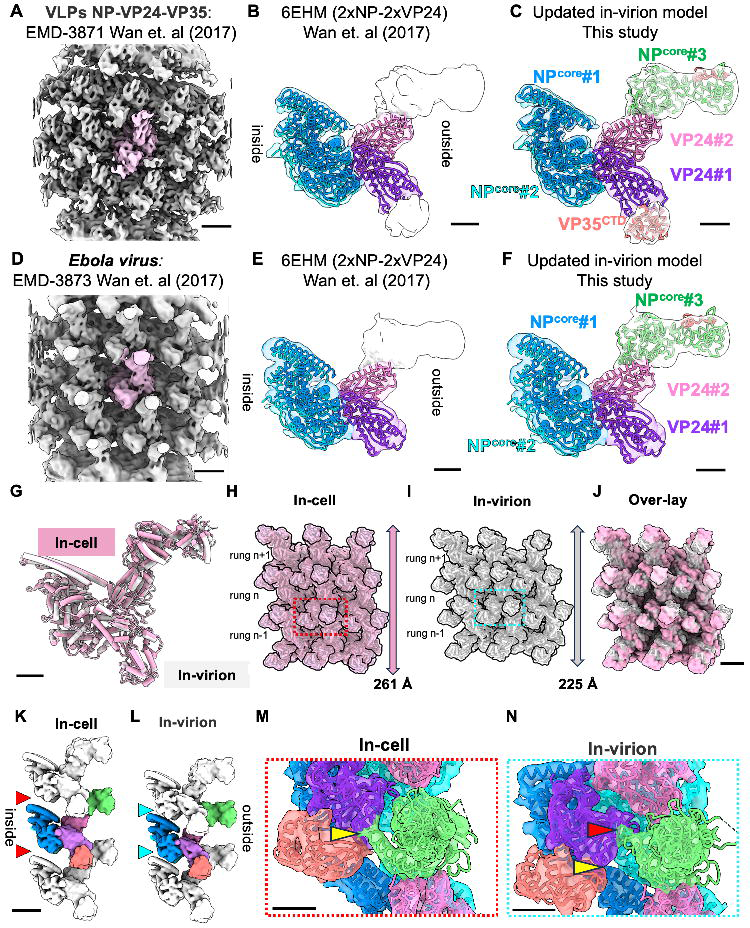
Molecular model of in-virion nucleocapsids and condensation of intracellular nucleocapsids upon virion incorporation. (A and D) Previously published EM maps of in-virion NP-VP24-VP35 (EMD-3871) and Ebola nucleocapsid (EMD-3873)^25^. The single repeating units are highlighted in pink. (B and E), EM densities fitted with the published molecular model of two copies of NP (in blue and cyan) and two copies of VP24 (in purple and pink) (PDB: 6EHM)^25^ molecules show previously unannotated densities (white). (C and F) Updated in-virion model created based on an in-cell model (Figure 2) fitted into EM densities of the virus like particles (VLPs) and authentic virus. Model colors correspond to those for Figure 2. (G) Comparison of single repeating units of intracellular (in-cell in pink) and in-virion (in white) nucleocapsid shows that the models are nearly identical. (H-J) Assembled models consisting of nine repeating units of in-cell and in-virion nucleocapsid models based on intracellular NP(Δ601-739)-VP24-VP35 and in-virion NP-VP24-VP35 nucleocapsid densities (EMD-3871)^25^ and their overlaid models. In the overlaid model, the in-cell surface model is shown in opaque pink. (K-L) Side view of assembled models consisting of three vertically aligned nucleocapsid models based on EM densities of intracellular NPΔ601-739-VP24-VP35 and in-virion NP-VP24-VP35 nucleocapsids(EMD-3871)^25^, respectively. Red and blue arrowheads show longer and shorter inter-rung gaps, respectively. (M-N) Enlarged views of the boxed regions in (H) and (I), respectively. Molecular models are fitted into the EM densities. Yellow arrowhead in (M) and red arrowhead in (N) indicate the inter-rung interfaces identified in the in-cell nucleocapsid and in the in-virion nucleocapsid, respectively. In (N), yellow arrowhead points to the same location of VP24 (purple) as in (M) as a reference point. Scale bars indicate 5 nm (A, D, J, and K) and 2 nm (B, C, E, F, G, M, and N). See also Figure S4 and Video S4.

Although the single repeating units are almost identical between in-cell and in-virion structures (Figure 4G), the inter-rung associations do substantially differ between in-cell and in-virion nucleocapsids (Figures 4H-4J and S4M-S4V). The helical pitches of intracellular nucleocapsids obtained here (85 Å and 87 Å for full-length NP and Δ601-739NP, respectively) are consistently larger than those previously reported for in-virion nucleocapsids^25^ (75 Å and 74 Å for VLPs and budded authentic Ebola virus, respectively). We propose that this longer inter-rung gap, rather than a different size of their repeating units, accounts for differences in their pitches (Figures 4K, 4L, and S4S-S4V, red and cyan triangles and Video S4). These data suggest that intracellular Ebola nucleocapsids are further condensed upon incorporation into virions.

### EBOV-GFP-**Δ**VP30-infected cells illuminate the nucleocapsid assembly process in the context of viral infection

We next visualized the nucleocapsid assembly process in the context of infection using a more authentic virus. To operate outside Biosafety level (BSL) 4 containment, we used a biologically contained model Ebola virus, EBOV-GFP-ΔVP30 (Figures 5, and S5). In this model, the gene encoding VP30, which is essential for viral transcription, is replaced in the viral genome by a gene encoding green fluorescence protein (GFP). EBOV-GFP-ΔVP30 virions can enter cells, but can not replicate unless VP30 is supplied in trans^34,35^. The GFP reporter indicates the presence of viral replication activity in infected cells that stably express VP30 (Figures S5A-S5C). In EBOV-GFP-ΔVP30-infected cells, similar to authentic virus infection, NP assembles on the viral RNA genome in the presence of all viral proteins, which, except for VP30, are all expressed under intrinsic control from the viral genome while VP30 is expressed by an exogenous promoter in the stable cell lines^34^. In this system, both virus and infected cells are operable outside a BSL4 environment.

**Figure 5.**
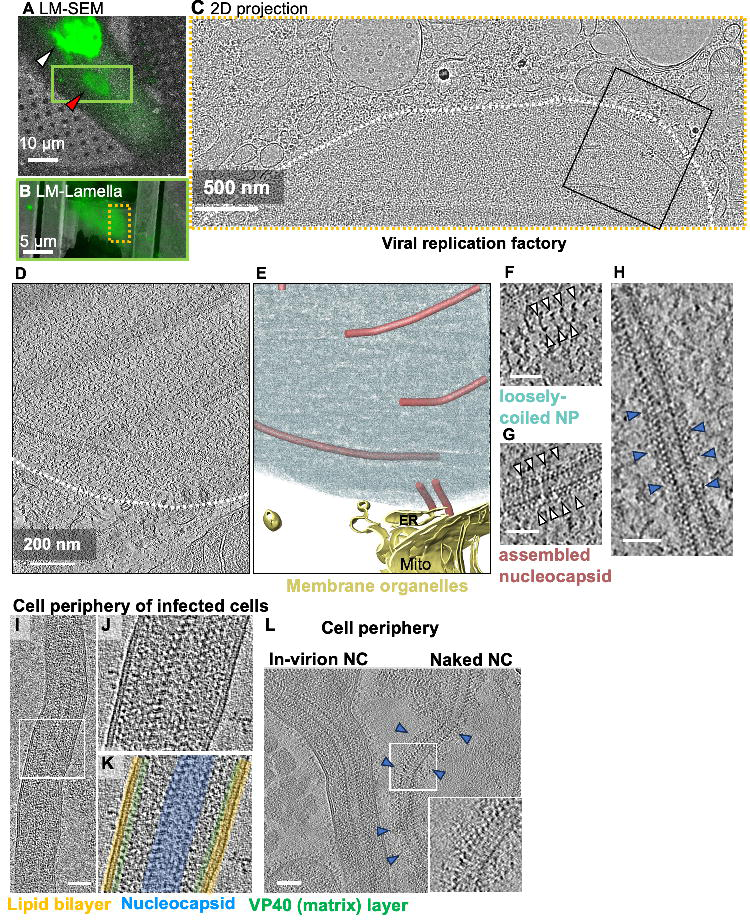
EBOV-GFP-ΔVP30 virus-infected cells illuminate the authentic assembly process. (A) Overlay of the fluorescence signal (green) and cryo-SEM image (greyscale) of chemically fixed and plunge-frozen EBOV-GFP-ΔVP30-infected VP30-Vero cell 24 hours post-infection. The locations of the nucleus and viral replication factory are marked in white and red arrowheads, respectively. (B) Overlay of the fluorescence signal (green) and an SEM image after FIB-milling of the area marked in (A). (C) TEM image of the area marked in (B) showing presence of virus replication factory in the host cell cytoplasm (marked with white dotted curve). (D) Tomographic slice of the area marked in (C). The replication factory is inside and above the white dotted curve. (E) Annotation of fully assembled nucleocapsid (in pink) and membrane (in yellow) and densities inside of viral factory (in blue gray) in the tomogram shown in (D). (F, G) The enlarged views of loosely coiled NP and fully assembled nucleocapsid, respectively. (H) Representative tomographic slice showing intracellular nucleocapsid found in EBOV-GFP-ΔVP30 virus-infected cells. Blue arrowheads indicate fluffy densities extending from the intracellular nucleocapsid. (I) Representative tomographic slice of a virus emerging from of chemically fixed and plunge-frozen EBOV-GFP-ΔVP30-infected VP30-Vero cell 24 hours post-infection. (J) Enlarged view of the boxed region in (I). (K) Annotation of tomogram in (J). The viral envelope lipid bilayer, matrix protein VP40 layer, and nucleocapsids are in yellow, green, and blue, respectively. (L) Representative tomographic slice of intact virus and naked nucleocapsids without viral envelope found in the cell periphery of infected cells 24 hours post-infection. Blue arrowheads highlight fluffy densities extending from the in-virion nucleocapsid. Inset: Enlarged views of the boxed region. Scale bars indicate 10 µm (A), 5 µm (B), 500 nm (C), 200 nm (D) and 50 nm (F-I and L). See also Figures S5.

During filovirus replication, nucleocapsid assembly and genome transcription and replication exclusively occur inside viral replication factories, also known as viral inclusion bodies, in the cytoplasm of the infected cell^33,36^. GFP signals were enriched in the viral factory and nucleus in chemically fixed cells (Figure 5A, S5C-S5E), but not in live infected cells (data not shown). This finding is likely due to the physicochemical properties of these compartments, which retain soluble GFP molecules through covalent chemical fixation^37^. We verified that VP30 and VP40 in addition to NP, VP24, VP35 (not shown) accumulated in replication factories (Figures S5D-S5E) as reported previously^38^. Using cryo-electron tomography coupled with cryo-fluorescence microscopy and focused-ion beam milling of plunge-frozen PFA-fixed EBOV-GFP-ΔVP30-infected cells, we observed intracellular virus replication 24 hours post-infection (Figure 5A-5H). We noted that most host membrane structures are often segregated from these virus replication factories (Figures 5C-5D, and S1P). The virus replication factory is filled with a loosely coiled NP, which closely resembles isolated full-length NP associating with cellular RNA^24^ and fully assembled nucleocapsid (Figures 5D-5G). We have noticed two differences between nucleocapsid structures from transfected and infected cells. First, in the transfected cells, we observe loosely-coiled, regionally condensed, and fully assembled nucleocapsid structures (Figure 1), whereas in virus-infected cells, we observe only loosely-coiled NP or fully assembled structures (Figure 5). Second, additional visible density protruding from the nucleocapsid is visible only in infected cells (Figure. 5H, highlighted with blue arrowheads). These differences are likely because authentic virus replication involves expression of all viral proteins and the nucleocapsids are destined for incorporation into the virus envelope at the cell surface (see discussion below).

We also visualized assembled EBOV-GFP-ΔVP30 virions emerging from infected cells by imaging the cell periphery of frozen-hydrated PFA-fixed EBOV-GFP-ΔVP30-infected cells using cryo-electron-tomography combined with cryo-fluorescence microscopy (Figures. 5I-5L, and S5F-S5G). In these emerging virions, the viral nucleocapsid lies at the center of the filamentous virion, and we visualize VP40 underlying the viral envelope, as previously reported with virus-like particles and authentic Ebola virus^39^. Like intracellular nucleocapsids in virus-infected cells, in-virion nucleocapsids in EBOV-GFP-ΔVP30 viruses are also surrounded by additional densities. Such structures of the budded nucleocapsids are particularly visible in particles that have lost the surrounding membrane envelope (Figure 5L).

### Conserved inter-molecular interfaces in filovirus nucleocapsid assembly

We suspect interfaces involved in the Ebola nucleocapsid assembly are essential and therefore well-conserved in the filovirus family. To test this hypothesis, we first identified interfaces using our intracellular model and map. Interfaces were identified by visually examining the atomic model fitted in the experimental map using the following criteria: (1) densities corresponding to the specified loops or helices are visible in the EM map; and (2), even increasing the threshold of the density, the connecting density between two proteins is still visible, assuming the regions forming interfaces are more rigid than elsewhere. We did not consider a region as an interface if the corresponding experimental EM densities were not observed, even if the distance shown in the model was small. We also observed potential interfaces where experimental density showed connections, but no specific residues were assigned in the atomic model.

Our molecular model of an intracellular nucleocapsid revealed at least six interfaces within the repeating unit, plus one interface that forms an inter-rung association between adjacent repeating units (Figure 6, and Table S1). At the current 9 Å resolution of our EM map, we can identify protein regions, but not the individual contact amino acids involved in each interface. Each of these interfaces is described in the order protein A-protein B in which protein A is toward the nucleocapsid interior, while protein B is toward the nucleocapsid exterior.

**Figure 6.**
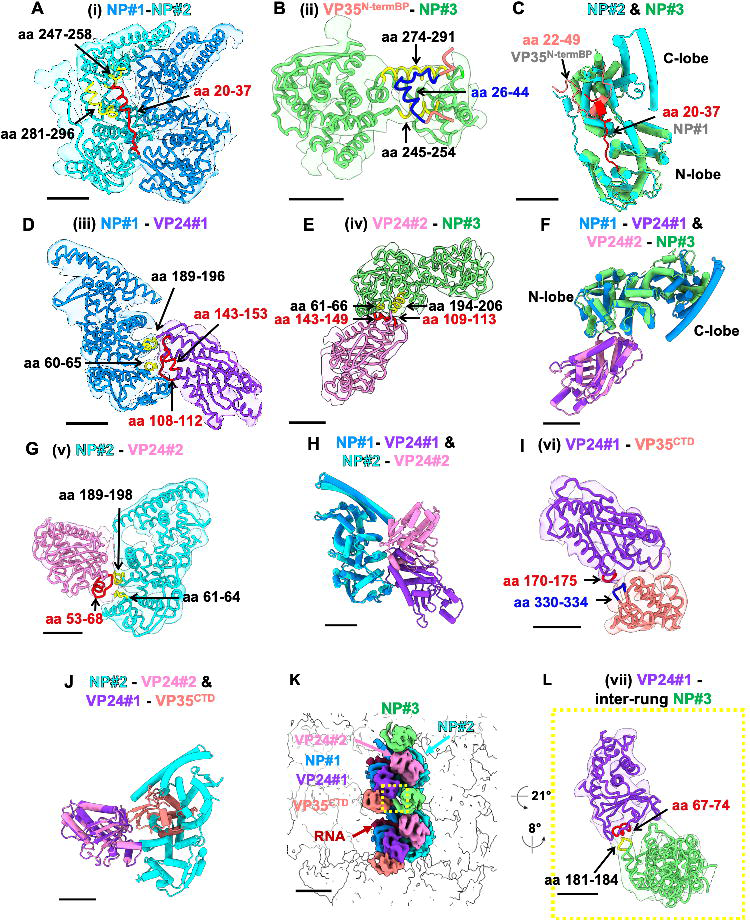
Interfaces involved in nucleocapsid formation. (A) Interface (i) NP#1-NP#2. The view is from inside the nucleocapsid as shown in Figure 2H. (B) Interface (ii) VP35^N-termBP^-NP#3. The view is from the top of the nucleocapsid as shown in Figure 2K. (C) Superimposing NP#2 and NP#3. N-terminal tail from the adjacent NP#1 and VP35 N-terminal binding peptide are shown in red and orange, respectively. (D) Interface (iii) NP#1-VP24#1. Side view as shown in Figure 2G. (E) Interface (iv) VP24#2-NP#3. Side view as shown in Figure 2G. (F) Superimposing NP#1-VP24#1 and VP24#2-NP#3 with alignment of the VP24#1 and VP24#2. (G) Interface (v) NP#2-VP24#2. Side view as shown in Figure 2I. (H) Superimposing NP#1-VP24#1 and NP#2-VP24#2 with alignment of NP#1 and NP#2. (I) Interface (vi) VP24#1-VP35^CTD^. Side view as shown in Figure 2G. (J) Superimposing NP#2-VP24#2 and VP24#1-VP35^CTD^ with alignment of VP24#2 and VP24#1. (K) Two central repeating units are colored within the EM density of intracellular NPΔ601-739-VP24-VP35. Yellow box shows an inter-rung interface. (L) Interface (vii) VP24#1-inter-rung NP#3. (A-L) The color of each fit model matches that of the EM map in Figure 2. Regions forming interfaces are colored yellow, red, and blue to distinguish them from the rest of the proteins. Density thresholds are set to show helical densities. Scale bars indicate 2 nm except in (K); 5nm. See also Figure S6 and Table S1.

The following features characterize the different interfaces: **(i)** The interface between NP#1 and NP#2, localized in the central NP-RNA helix, is mediated by insertion of the N-terminal helix (aa 20-37, red) of NP#1 into the pockets of the adjacent NP#2 that are formed by aa 247-258 and 281-296 [yellow; (Figure 6A)]. This interface is identical to that observed in assembled recombinant NP N-terminal domain (1-450) and in nucleocapsids of budded virions^25,40^.

**(ii)** The interface between VP35^N-termBP^ and NP^core^ of NP#3, on the outer layer of the nucleocapsid, is mediated by an interaction between the N-terminal NP-binding peptide of VP35 (NP interaction site is aa 26-44 of VP35 in blue, bracketed by aa 22-25 and 45-49 of VP35 in orange) with the C-terminal lobe of the NP core (aa 245-254 plus 274-291 of NP, yellow; Figure 6B) and is identical to that seen in VP35-NP crystal structures^11,12^. Superimposing NP#2 and NP#3 confirms that the N-terminal helix of NP#1 and the N-terminal NP-binding peptide of VP35 both insert into the same narrow pocket created by the two alpha helices of NP, indicating a single NP can bind only one of these binding partners at a time and that what partner is bound there determines whether NP is in a monomeric or an oligomeric state (Figure 6C). We also note that arrangements of part of the penultimate and long clamp helices (aa 355-405) in the C-lobe differ between NP#2 and NP#3 (Figure 6C, 2J and 2K).

**(iii)** The interface of NP#1-VP24#1 is mainly mediated by two loops of the NP (aa 60-65 and 189-196, yellow; Figure 6D) and two loops of VP24 (aa 108-112 and 143-153, red; Figure 6D). **(iv)** Interestingly, almost the same interfaces are used by the interaction of VP24#2 (aa 109-113, 143-149, red) with the outer NP#3 (aa 61-66, 194-206, yellow). Superimposing NP#1-VP24#1 and VP24#2-NP#3 further validates that the same VP24 and NP interaction occurs at two different locations within the repeating unit, one facing inward and the other facing outward in the nucleocapsid (see above, Figure 6F).

**(v)** In the interface between NP#2 and VP24#2, a VP24 loop that includes aa 53-68 (red; Figure 6G) interacts with two NP loops (aa 61-64, 189-198; yellow Figure 6G). Alanine substitution of aa 53-59 of VP24 was previously shown to abrogate NP binding in co-immunoprecipitation experiments performed in the absence of VP35^16^, providing biochemical evidence for interface (**v)**. Superimposing the NP#1-VP24#1 and NP#2-VP24#2 complexes reveal that nearly identical NP surfaces are involved in interactions, but the two bound VP24 are in opposite orientations (Figure 6H).

**(vi)** The interface of VP24#1-VP35^CTD^ is mediated by loops from VP24 (aa 170-175, red; Figure 6I) and VP35 (aa 330-334, blue; Figure 6I). Superimposing NP#2-VP24#2 and VP24#1-VP35^CTD^ reveals that VP35^CTD^ and NP bind to the same side of VP24: the binding of VP35^CTD^ to VP24#1 overlaps with the binding of NP#2 to VP24#2 (Figure 6J). The overlap of binding suggests that binding of VP35^CTD^ to VP24#1 prevents a second NP from binding this VP24#1.

**(vii)** The interface between VP24#1 with NP#3 on the next rung down (yellow box, Figure 6K) mediates inter-rung interactions of the intracellular nucleocapsid via a helix from VP24 (aa 67-74, red) and a loop from NP (aa 181-184, yellow; Figure 6L). Alanine substitution of aa 73-77 of VP24 was previously shown to abrogate NP binding in co-immunoprecipitation experiments performed in the absence of VP35^16^, providing biochemical evidence for this newly visualized VP24#1-inter-rung NP#3 interface (**vii)**.

The inter-rung distances differ significantly between in-cell and in-virion nucleocapsids (Figures 4H-4L, S4M-S4V, Video S4). In-cell nucleocapsids are vertically more open, whereas in-virion nucleocapsids are more condensed. A side-to-side comparison of the corresponding regions reveals that the inter-rung interface (**vii)** in the in-cell nucleocapsid (Figure 4M, yellow arrowhead) is not present in the in-virion nucleocapsid. In the in-virion nucleocapsid, the potential inter-rung interface between VP24#1 and NP#3 on the next rung down is shifted upward (Figure 4N, red arrowhead. The yellow arrowhead points to the same position as Figure 4M as a reference point). At the resolution of the existing published map of the nucleocapsid in virus-like particles (EMD-3871^25^), we cannot define the presence or identity of potential inter-rung interactions in the virion.

We next examined if the regions comprising the seven interfaces (**i** to **vii**) are well-conserved among six different filoviruses (Figure S6). After sequence alignment with T-Coffee (Figures S6A-S6C), we calculated the average conservation score of these interface regions using three different metrics. We compared them to scores for the full-length proteins (Figure S6D and Table S1). All three metrics indicated that the conservation scores for all the interface regions are substantially higher than the average score for the full-length proteins, suggesting that nucleocapsid assembly architecture and the interfaces are likely conserved across different filoviruses.

## Discussion

In this study, we sought to understand the assembly process of the Ebola virus nucleocapsid in producer cells, prior to viral egress. We visualize not only fully assembled nucleocapsids, but also identify distinct nucleocapsid assembly intermediates at different states of condensation (Figures 1, 5 and 7). This analysis further allowed identification of a previously unknown third layer of the Ebola virus nucleocapsid, which includes NP maintained in a monomeric, genome-free state by VP35 (Figure 2 and Videos S1-S3), as well as the likely location of the NP C-terminal domain (aa 601-739) at the outside of the nucleocapsid, deduced by visualization of how nucleocapsids bundle in transfected cells (Figure 3).

**Figure 7.**
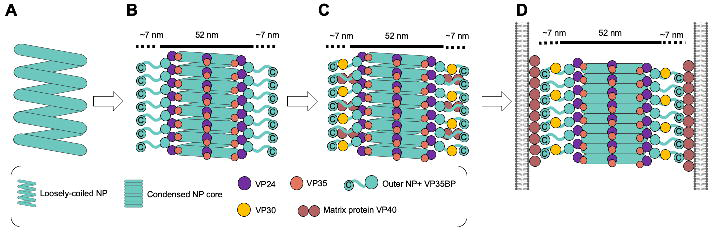
Schematic model of the Ebola virus nucleocapsid assembly process. (A) The loosely coiled NP. (B) VP24, VP35, and outer NP assemble on the loosely coiled NP, which transition into a rigid and condensed state. A portion of the intrinsically disordered domain and C-terminal domain of NP extends from the assembled nucleocapsid and fills the additional 7 nm-wide space along the intracellular nucleocapsid. (C) In virus-infected cells, the essential transcription factor VP30 and matrix protein VP40 interact with C-terminal domains of NP^46–48,52,53^. (D) Assembled nucleocapsids are incorporated into virions. In the virion, nucleocapsids are further condensed relative to intracellular nucleocapsids. The matrix-to-matrix distance in the virion measured in the tomographic slice shown in Figures 5I is ∼66 nm, which matches well with the estimated width of ∼66 nm of nucleocapsid including NP (aa 601-739) regions (52 + 14 nm) shown in Figures 3F and 3I.

Our surprising finding of a monomeric NP molecule in assembled nucleocapsid is the first example among *Mononegavirales.* We now see that in the Ebola virus nucleocapsid, both polymerized RNA-bound and monomeric RNA-free NP exist. The outer NP is held monomeric and RNA-free by binding to the phosphoprotein VP35. We propose that this unexpected third NP functions to efficiently link the nucleocapsid to the viral matrix protein VP40 to incorporate the nucleocapsid into the virus particles in the budding process. Both the N-terminus (aa 26-150) and C-terminal domain of NP (601-739) are required for matrix protein interactions and nucleocapsid incorporation into the virion^46–48^.

In both Ebola virus and Marburg virus, the NP-RNA central helix is surrounded by a 6-7 nm thick layer of repeated copies of VP24^25^. VP24 is unique to the filoviruses and is known to play a role in host immune evasion^18–21,49^. Thus, carrying multiple copies of VP24 (two VP24 for every three NP) in the viral payload may limit host immune responses from the earliest time points after infection. However, this 7 nm thick layer of VP24 envelops the central NP helix and could hinder interactions between NP and the globular VP40 dimers that would be needed to incorporate the nucleocapsid into the virion^39,50^. In particular, the NP N-terminus (aa 26-150), essential for VP40 interaction, is buried in the central NP-RNA helix in the inner-layer copies of NP. In contrast, in the NP on the outer layer, the N-terminus is fully exposed and easily accessible for VP40 matrix interactions. We propose that the different layers of NP assembled in the Ebola nucleocapsid are tuned to perform two distinct functions. One function, achieved by NP#1 and NP#2 in the core, is to bind RNA and polymerize to form the central NP-RNA helix and protect the viral genome. The other function, achieved by NP#3, is to interact with the matrix proteins to drive nucleocapsid incorporation into the virion.

This work revealed that VP35 interacts with the outer layer of NP and predicted a location of VP35^CTD^ (aa 221-340) in association with VP24#1. We attempted to fit a more complete structure of VP35, amino acids 81-340, the longest VP35 structure experimentally determined (PDB: 7YER^15^ (Figure S7), into the nucleocapsid model based on the location of VP35^CTD^. In this speculative model, the central oligomerization domain of VP35 (aa 81-145) points toward NP^core^#3 of the adjacent repeating unit. Part of the VP35 oligomerization domain (aa. 81-112) may interact with the outer NP molecule. Although not yet experimentally visualized, this hypothetical second interaction between the outer NP and VP35 agrees with previously published biochemical immunoprecipitation assays^11^. Previous work identified that VP35 contains two sites critical for NP binding: the N-terminus that chaperones NP into a monomeric and RNA-free state (aa 1-80), as well as a second, independently functioning NP binding site (aa 80-120) in the oligomerization domain^11^. The previously demonstrated flexibility of the VP35 oligomerization domain (aa 81-145)^51^ might explain the lack of corresponding densities for this VP35 region in our EM map. The bridge-like conformation of the extended VP35 molecule might suggest that the outer NP^core^#3 takes part in an intra-rung nucleocapsid interaction via interaction with VP35 (Figure S7). This finding is consistent with the requirement for VP35 in the formation of the outer layer of the nucleocapsid that lies outside the core NP^core^-RNA helix^5,6^.

We also reveal structural rearrangements between in-virion and in-cell nucleocapsids. The in-cell nucleocapsids have a more open structure and further, additional inter-rung interactions are only visible in-cell. These additional inter-rung interactions reveal how the viral nucleocapsid condenses vertically as the nucleocapsid matures in its final budded in-virion form (Figure 4, 7 and Video S4). Interestingly, in our 3D classification of intracellular nucleocapsids, some regions exhibit shorter inter-rung distances even within the same particles (Figures S2W). These more condensed regions have narrower inter-rung distances and lack the outer NP-VP35 complex (Figure S2W, Classes 6 and 9 marked in red arrowhead outline). These findings imply that the outer NP-VP35 complex bridges interactions between adjacent repeating units that may determine some optimal inter-rung distance of an intracellular nucleocapsid. Vertical condensation of the nucleocapsid (Figures 4, 7C-7D and Video S4) may protect RNA genome integrity through tight NP-inter-rung associations mediated by electrostatic interactions^25,40^. Condensation may be triggered by interaction of the outer NP with matrix protein along the viral membrane upon incorporation into virion.

It was previously described using recombinant protein that the NP N-terminal domain (aa 1-450), but not full-length NP, can form condensed helices encapsidating cellular RNA with dimensions and pitch similar to the inner nucleocapsid core in the virion^24,25,40^. These results suggest that the C-terminal half of NP (aa 451-739) could hinder this tight condensation. One of our observations here, comparing nucleocapsid structures between transfected cells (NP+VP24+VP35) and infected cells (EBOV-GFP-ΔVP30; all genes present), is that in infected cells, we only observe either loosely coiled NP or fully assembled nucleocapsids (observed with additional outer density). Whereas in the NP+VP24+VP35 transfected cells, we observed loosely-coiled NP, fully assembled nucleocapsid, and partially/regionally condensed NP (Figures 1 and 4). The spontaneous regional condensation of NP seen in transfected cells (Figure 1) suggests that full-length NP is capable of a tightly condensed helical structure, albeit at a much lower efficiency than the NP N-terminal domain alone. However, in virus-infected cells, NP preferentially associates with the viral RNA genome (absent in transfected cells), uses its C-terminus (aa 601-617) to bind VP30^52,53^ and recruits RNA-dependent RNA polymerase L via interaction with VP35 to facilitate RNA genome replication and viral mRNA transcription^54^. We suspect that all these activities, and the proteins associated with NP to achieve them, together prevent spontaneous regional condensation of NP. In virus-infected cells, recruitment and physical binding of VP24, VP35, and outer NP might be strictly required to condense the NP helix. This condensation is likely critical to shift the transcriptionally active, more open nucleocapsid into its inactive, condensed structure before incorporation into the virion (Figure 7).

We observed that both in-cell and in-virion nucleocapsids, each expressed with all viral genes present, showed additional densities emerging from nucleocapsids (Figure 5) that are not visible when only NP, VP24 and VP35 are transfected. The additional densities may be unknown host proteins or other virus proteins such as VP30 and VP40, both of which have been shown to interact with the C-terminal region of NP (aa 601-749)^46–48,52,53^. Both VP30 and VP40 localize in viral replication factories in EBOV-GFP-ΔVP30-infected cells (Figure S5). For both Ebola virus and Marburg virus, the viral nucleocapsid lies at the center of the filamentous virion, surrounded by VP40 matrix proteins^39^ (Figure 5). The existence of some hypothetical flexible linker-like structure, of unknown identity, connecting the nucleocapsid to the matrix has been proposed to minimize nucleocapsid breakage when the long filamentous filoviruses (1 to 2 µm) bend^55^. In the tomographic slice shown in Figure 5I, the VP40-to-VP40 distance in-virion is roughly 66 nm, which matches well with the predicated 66 nm width of the entire nucleocapsid (52 nm) that includes the NP C-terminal domain (7 nm both sides, total 14 nm) (Figures 3F). These observations support the idea that in infected cells, the NP C-terminal region (aa 601-739) recruits other viral proteins to the nucleocapsids and functions as a flexible linker between the nucleocapsid and matrix proteins in the virion (Figure 7) to protect viral genome integrity.

### Limitations of the study

Our *in situ* structural analysis of the fully assembled Ebola virus nucleocapsid revealed the previously unknown complex organization of intracellular nucleocapsid structures and revealed the numbers of intermolecular interactions between the separate copies of NP, VP24 and VP35 that are conserved among distinct filovirus species. In this study, we could not assign the precise amino acid residues involved in these interfaces due to the current resolution of our EM map. While this manuscript was under submission, *in situ* cryo-ET study of intracellular nucleocapsid assembly process in authentic Ebola virus focusing on morphological changes of nucleocapsid structures in viral replication factories during viral infection progress has been reported^56^. The authors reveal that Ebola viral nucleocapsids undergo a transition from loosely coiled helical assemblies to condensed cylinders organized into parallel bundles in viral replication factories as infection progresses, agreeing with our claim that loosely-coiled NP is the precursor of fully assembled nucleocapsid during viral assembly. Another cryo-EM single particle study of Ebola nucleocapsid in viral-like particles reported an improvement in resolution (4.6 Å) of the previously defined inner and middle layer of the in-virion Ebola nucleocapsid and identified critical residues forming those interfaces^57^. However, in both studies, the outermost layer newly assigned as 3rd NP and VP35 in this study was not revealed, likely due to the flexible nature of the outer layer. In addition, there is no structural information of the filovirus-specific, long, intrinsically disordered C-terminal region of NP (aa 405-600) in any filovirus nucleocapsid structure. The region of NP (aa 451-600) is critical for nucleocapsid formation in the presence of VP24 and VP35^26^ and therefore, more studies will be required to illuminate this portion of the Ebola virus nucleocapsid assembly.

## Author contributions

R.W. conceived the project and designed the experiments under the supervision of E.O.S. D.P. created plasmid encoding NPΔ601-739. Y.J. and G.C. prepared resin-embedded samples. R.W. prepared the samples, performed fluorescence microscopy, performed cryo-FIB milling, acquired the cryo-electron tomography data, and performed image processing with help from D.Z., C.H., and W.W. D.Z. and R.W. performed model building. R.W., S.L.S and E.O.S. analyzed and interpreted the data and wrote the manuscript with the help of comments from all authors.

## Acknowledgments

We thank Drs. Digvijay Singh (UCSD), Vinson Lam (University of Michigan), Thomas G. Laughlin (Altos Lab), Benjamin Barad (TSRI), Hamidreza Rahmani (TSRI), Jingru Fang (LJI), Sara Landeras Bueno (LJI), Theresa Gewering (LJI), Lea Dietrich (Max Planck Institute of Biophysics), Ben Engel (Biozentrum, Basel) Ricardo Diogo Righetto (Biozentrum, Basel) and members of the Saphire lab for providing technical and biological expertise throughout the project. Grid preparation, cryogenic-fluorescence microscopy, cryo-focused ion beam milling, and cryo-electron tomography were done at the LJI cryoEM core. We thank Dr. Ruben Diaz Avalos for discussions on data acquisition and processing as well as management of the instruments. Resin-embedded TEM sample preparation and data acquisition was performed at the Cellular and Molecular Medicine Electron Microscopy Core (RRID: SCR_022039). Room temperature fluorescence microscopy was performed at the LJI Microscopy and Histology core. We thank Drs Zbigniew Mikulski and Sara McArdle for training and discussion. We thank Drs. Yoshihiro Kawaoka and Peter Halfmann (University of Wisconsin) for providing the anti-VP30 and anti-VP24 antibodies, the EBOV-GFP-ΔVP30 system, as well as sharing their expertise, and Dr. Laurence Cagnon (LJI Biosafety officer) for instruction in handling EBOV-GFP-ΔVP30 system under BSL2 laboratory with BSL3 practices. Drs. Jason Greenbaum and Ashmitaa Logandha Ramamoorthy Premlal of the LJI Bioinformatics core calculated the conservation score of filovirus orthologs; we thank them for their guidance on the conservation analysis. We thank Drs. Michael Scarpelli, Michael Talbott, and their team at the LJI IT Infrastructure and HPC Technologies for technical support.

## Supplemental figure legends

**Figure S1. NP and NPΔ601-739 colocalize with VP24 and VP35 and they form nucleocapsid-like structures in cells indistinguishable from EBOV-GFP-ΔVP30 virus-infected cells, related to Figures 1, 2, and 3**

(A-C) Fluorescence microscopy images of HEK 293T cells transiently expressing NP, VP24, and VP35

(D-F) Fluorescence microscopy images of HEK 293T cells transiently expressing NPΔ601-739, VP24, and VP35

(A-F) Cells were fixed with 100% Methanol at −20 °C and co-stained with rabbit polyclonal anti-NP and mouse monoclonal anti-VP35 antibodies (A and D), rabbit polyclonal anti-NP and mouse monoclonal anti-VP24 antibodies (B and E), and rabbit polyclonal anti-VP35 and mouse monoclonal anti-VP24 antibodies (C and F). The nuclei were stained with Hoechst. The upper two panels showed images for individual channels and the bottom panels showed overlaid images. The presented images showcase representative images derived from three independent experiments.

(G-P) Ultra-thin resin-embedded transmission microscopy images of HEK 293T cells transiently expressing NP, VP24, and VP35 (G), NPΔ601-739, VP24, and VP35 (H), NP alone (I), NPΔ601-739 alone (J), VP35 alone (K), VP24 alone (L), VP35 and VP24 (M), NP and VP24 (N), NP and VP35 (O) or EBOV-GFP-ΔVP30-infected VP30-Vero cells (P). An expanded view of the area enclosed by the box on the left panel is shown in the right panel. The blue arrowheads indicate fully assembled, condensed nucleocapsids.

Scale bars represent 15 µm (A-F) and 2 µm (G-P).

**Figure S2. Subtomogram averaging, related to Figures 2 and 3**

(A) Preprocessing and tomogram reconstruction were done in Warp based on the patch track alignment done with ETOMO. Nucleocapsids were traced using the Amira fiber tracing module with 4-bin deconvolved tomograms.

(B) The center line of filaments was equidistantly re-sampled (4 nm). After initial alignment in Dynamo using an 83 nm box size, we estimated helical parameters using HI3D.

(C-D) Re-cropping and initial averages of subtomogram centered on the nucleocapsid double layer in Dynamo (C) and in Warp (D).

(E-G) RELION 3D classification without alignment (E). 3D refinement with a soft mask covering the nucleocapsid with 2-bin (F) and unbinned data (G).

Class 1 and 2 particles from the initial RELION 3D classification (E) were further analyzed as shown in Figures S2W and S2X.

(H) Final sharpened map with M.

(I) Angular distribution plot of all particles that contributed to the final map. The height and color (from blue to red) of the cylinder bars is proportional to the number of particles in those views.

(J) Gold-standard Fourier Shell Correlation plots obtained with the RELION Post-processing module and estimated at 9.37 Å.

(K) Local resolution estimation obtained in M, and maps were rendered with ChimeraX.

(L-R) The same procedures carried out for NPΔ601-739 were done for full-length NP except that box sizes of 90 nm (M) and 43 nm (N) were used.

(S) Final sharpened map with M with 2-bin data.

(T) Angular distribution plot of all particles contributed to the final map.

(U) Gold-standard Fourier Shell Correlation plots were obtained with the RELION Post-processing module and estimated at 18 Å.

(V) Local resolution estimation obtained in M, and maps were rendered with ChimeraX.

(W) Re-classification of particles sorted to classes 1 and 2 from intracellular NPΔ601-739-VP24-VP35 nucleocapsid data (Figure S2E) into 10 classes (now termed classes 1-10). The squares outlined by dashed line highlight the regions of interest (the narrower inter-rung distance). The outline or filled red arrowheads point to the absence or presence, respectively, of outer NP within the same particles.

(X) The positions of the individual particles were plotted back to the representative tomogram using ArtiaX. The particle distribution in the tomogram was visualized with two angles (Upper panels: x-z, the lower panels; x-y views). Green, red, and gray particles correspond to the particles that contribute to the fully assembled nucleocapsid average, particles sorted into the indicated class, and all other remaining particles in the tomogram, respectively.

We note that Class 2 particles are exclusively sorted into both surfaces of the tomogram, suggesting they are incomplete particles and could have sustained damage during FIB milling^59^. All the other particles are scattered throughout the tomogram, and are even located next to particles from the best class, suggesting that intracellular nucleocapsids assemble into highly heterogeneous populations.

**Figure S3. Model building of intracellular Ebola virus nucleocapsid, related to Figure 2**

(A) The table summarizes previously published atomic models used in this study. Modeled regions are indicated by residue numbers.

(B) RNA density encapsidated in intracellular nucleocapsid is shown in dark red.

(C-J) The molecular models fitted into the EM densities for NP#1-NP#2 (C), VP24#1 (D), VP24#2 (E), VP35^N-termBP^-NP#3 (F and G), and VP35^CTD^ (H-J). Density thresholds are set to show helical densities.

Scale bars indicate 2 nm.

We note there is an unassigned density indicated with black arrows in (I-J). This density might correspond to the part of the unstructured C-terminal region of VP24 (aa 232-251), which was not modeled in pre-existing VP24 crystal structure PDB: 4M0Q^30^ used for our integrative modeling. The VP24 (aa 231) modeled in our molecular model, locates close to this density. However, due to the lack of sufficient resolution with our current EM map, we decided not to model this identity.

**Figure S4. Comparison of various nucleocapsid structures, related to Figures 3 and 4**

(A-L) The intracellular NP-VP24-VP35 (A, G) and NP(Δ601-739)-VP24-VP35 (C, I) nucleocapsid and overlaid images (B, H) and corresponding cross-sectional views (D-F, J-L).

(M-R) Comparison of EM maps for intracellular NPΔ601-739-VP24-VP35 nucleocapsid (M and P; pink) and in-virion NP-VP24-VP35 nucleocapsid (EMD-3781)^25^ (O and R; gray) and their superimposition (N and Q).

(S-V) Side view of assembled models showing three vertically aligned nucleocapsid models based on the maps of intracellular NPΔ601-739-VP24-VP35 (S, same as Figure 4K), intracellular NP-VP24-VP35 (T), in-virion NP-VP24-VP35 nucleocapsid (U, same as Figure 4L), and Ebola nucleocapsids (V). Red and blue arrowheads show longer and shorter inter-rung gaps, respectively.

Scale bars indicate 10 nm (A and C), 5 nm (G-R, and V).

**Figure S5. Visualization of EBOV-GFP-ΔVP30-infected cells, related to Figure 5**

(A) Genome organization of Ebola virus and biologically contained model Ebola virus in which the VP30 gene open reading frame is replaced by a gene encoding green fluorescence protein (GFP).

(B) Characterization of EBOV-GFP-ΔVP30. VP30-Vero cells (center, right panels) or wild-type Vero cells (left panels) were infected with EBOV-GFP-ΔVP30 (left and center panels) or mock-treated (right panels). 72 hours post-infection, cells were fixed with 4 % PFA and nuclei were stained with Hoechst. GFP signal indicates active viral replication (central upper panel). Nuclear staining shows that the integrity of the EBOV-GFP-ΔVP30-infected VP30-Vero cell monolayer is severely compromised due to viral replication (central lower panel) relative to the other conditions. The presented images showcase representative images derived from three independent experiments.

(C) Representative cryo-fluorescence microscopy image of chemically fixed and plunge-frozen EBOV-GFP-ΔVP30-infected VP30-Vero cells. GFP signals and fluorescence reflection imaging are in green and gray, respectively. The locations of the nucleus and viral replication factory are marked in white and red arrowheads, respectively.

(D-E) Immunofluorescence microscopy images of EBOV-GFP-ΔVP30-infected VP30-Vero cells fixed with 4 % PFA after 72 hours of infection and staining rabbit polyclonal anti-VP30 (D) or mouse monoclonal anti-VP40 (E). Yellow arrowheads indicate viral replication factories. VP30 was highly enriched in viral replication factories, while VP40 enrichment was observed in both the factories and at the plasma membrane. The presented images showcase representative images derived from four independent experiments.

(F) TEM image overlaid with GFP fluorescence image of EBOV-GFP-ΔVP30 virus-infected cells fixed and plunge-frozen 24 hours post-infection as visualized in Maps. Infected cells show plasma membrane blebbing (red arrowheads), suggesting the cells are in a late stage of apoptosis due to virus infection.

(G) Enlarged view of the boxed area shown in (F). A filamentous virus with an assembled nucleocapsid is visible. The tomogram taken here is shown in Figure 5I-K.

**Figure S6. Sequence alignment of orthologs of NP, VP24, and VP35 and regions involved in the interface and average conservation scores, related to Figure 6**

(A-C) Seven interfaces identified in Figure 6 are highlighted in corresponding color bars above the sequences of NP (A), VP24(B), and VP35 (C) for different filoviruses. *, : and . represent identical in all sequences in the alignment, conserved substitutions have been observed, and semi-conserved substitutions are observed, i.e., amino acids having similar shapes, respectively.

(D) Comparison of the conservation score averages for regions involved in interfaces i to vii (Figure 6) and across full-length proteins. Blue, purple, and orange dotted lines represent average scores across the entire amino acid sequence of NP, VP24, and VP35, respectively. Scores were calculated using the Karlin metric. The maximum conservation score, in which an amino acid sequence is identical across all 6 filoviruses, is 1 in this metric.

**Figure S7. Potential location of VP35 oligomerization domain, related to Figures 2**

(A-C) Three repeating units of in-cell nucleocapsid models are fitted in EM density of intracellular NPΔ601-739-VP24-VP35. The longest experimental VP35 structure (PDB: 7YER aa 81-340 in yellow)^15^ is fitted based on the VP35^CTD^ (aa 221-340) densities (orange) of the center and right repeating units.

(D) Potential interaction of outer NP#3 with the VP35 oligomerization domain. Outer EM density corresponding to the area shown as red box in (B). VP35 (aa 81 to 112 highlighted by transparent blue) could potentially interact with NP#3.

Scale bars indicate 2 nm

The color of each fit model matches that of the EM map in Figure 2.

## STAR Methods

### RESOURCE AVAILABILITY

#### Lead contact

Erica Ollmann Saphire;

#### Materials availability

Any requests for materials synthesized during this study should be addressed to Erica Ollmann Saphire.

#### Data and code availability

Maps have been deposited in the EMDB under accession codes EMD-42509 (intracellular nucleocapsid of NP601-739), and EMD-42515 (intracellular nucleocapsid of full-length NP). Models have been deposited in the PDB under accession codes 8USN (intracellular nucleocapsid model), and 8UST (in-viron nucleocapsid model).

This paper does not report original code.

### EXPERIMENTAL MODEL ANS SUBJECT DETAILS

#### Cell culture

Human embryonic kidney HEK 293T cells (ATCC reference: CRL-3216), African green monkey kidney tissue derived Vero cells (ATCC reference: CCL-81) were originally purchased from the ATCC. Vero cells stably expressing Ebola virus VP30 protein, VP30-Vero cell was obtained from Dr. Yoshihiro Kawaoka (University of Wisconsin, Madison). All cell lines were maintained in Dulbecco’s modified Eagle medium (DMEM-GlutaMAX) [Thermo Fisher Scientific, (TFS)], supplemented with 4.5 g/L D-Glucose,10% fetal bovine serum (FBS), penicillin (100 U/mL), streptomycin (100 μg/mL) in a 5 % CO_2_ incubator at 37 °C. Puromycin (5 μg/mL) was included in the media for culture of VP30-Vero cells but was excluded for the infection experiment. The biologically contained Ebola virus, EBOV-GFP-ΔVP30, was generated in HEK 293T cells and grown in VP30-Vero cells as previously described^34^. Virus was aliquoted and kept at −80 °C until use.

#### Plasmids

Plasmids encoding full-length Ebola virus NP and VP35, pCAGGS_NP_EBOV (#103049) and pCAGGS_VP35_EBOV(#103050), respectively, were gifts from Dr. Elke Mühlberger^60^ and purchased from Addgene. To create a plasmid encoding Ebola virus NP lacking amino acid 601-739, pCAGGS_NPΔ601-739_EBOV was created using NEBuilder HiFi DNA Assembly (NEB). After confirming the coding sequence, the fragment was subcloned into the original pCAGGS vector. The entire plasmid sequence was confirmed by Priomordium. pCAGGS_VP24_EBOV and all plasmids required to produce the biologically contained Ebola virus, EBOV-GFP-ΔVP30, were obtained from Dr. Yoshihiro Kawaoka (University of Wisconsin, Madison).

### METHODS DETAILS

#### Transfection

HEK 293T cells were transfected with pCAGGS_NP_EBOV or pCAGGS_NPΔ601-739_EBOV together with pCAGGS_VP35_EBOV and pCAGGS_VP24_EBOV using Lipofectamine 3000 (TFS) according to the manufacturer’s protocol.

#### Biosafety level of biologically contained EBOV-GFP-**Δ**VP30

La Jolla Institute for Immunology has obtained NIH approval to work on EBOV-GFP-ΔVP30 at Biosafety Level (BSL) 2 physical containment with selected BSL3 practices and enhanced biosecurity, so-called BSL2 plus. Before leaving this entry-controlled designated area, infected cells cultured on plastic dishes, cover glasses or TEM grids, and materials exposed to infected cells and media were chemically inactivated with 2.5 % glutaraldehyde at 4 degrees for 1 hour (ultra-thin resin-embedded transmission electron microscopy) or 4 % paraformaldehyde at room temperature for 15 min (immunofluorescence microscopy and cryo-FIB-ET). La Jolla Institute for Immunology Institutional Biosafety Committee approves these chemical inactivation procedures to downgrade samples as BSL1 materials and allows all following procedures to be performed at BSL1 containment.

#### Virus infection with EBOV-GFP-**Δ**VP30

VP30-Vero cells were seeded day before infection. The next day, the cells were infected with EBOV-GFP-ΔVP30 (MOI = 3 foci-forming unit/cell) by exchanging media with virus-containing supernatant and incubating for 1 hour in a CO_2_ incubator with gentle swirling the media every 10 min. After 1 hour, the virus supernatant was removed, and the cells were washed once with warm DMEM-GlutaMAX supplemented with 4.5 g/L D-Glucose, 2% FBS, penicillin (100 U/mL), streptomycin (100 μg/mL) and then incubated in the same media for the indicated time in a CO_2_ incubator. To terminate infection and inactivate the virus, infected cells were treated with 4% PFA in 0.10 M sodium cacodylate buffer pH 7.4 for 15 min.

#### Fluorescence microscopy of fixed transfected or infected cells

HEK 293T cells (2-6 x 10^5^ cells/well) and VP30-Vero cells (1-2 x 10^5^ cells/well) were plated onto 6-well dishes containing acid-washed, poly-L-lysine-coated (Sigma #P4707) and acid-washed (without coating) coverslips the day before the transfection and infection, respectivey. After indicated time, transfected or infected cells were fixed with 100% Methanol at −20 °C or 4% paraformaldehyde (PFA) in 0.1 M HEPES pH. 7.4 for 15 minutes at room temperature, respectively. Cells were washed three times in phosphate-buffered saline (PBS) and processed for antibody staining after blocking with 3% bovine serum albumin in PBS (Sigma #7906) to minimize non-specific binding. Antibody staining was done in the presence of 1 % BSA in PBS for at least 1 hour at room temperature. Anti-NP rabbit polyclonal (GeneTex, #GTX134031, 1: 300), anti-VP24 mouse monoclonal (Kawaoka lab, clone mouse/21.5.2.5, 1:2000), anti-VP35 rabbit polyclonal (GeneTex, #GTX134032, 1:300) or anti-VP35 mouse monoclonal (Kerafast, #EMS702 clone mouse/6C5, 1:300), anti-VP30 rabbit polyclonal (Kawaoka lab, 1:2000), anti-VP40 mouse monoclonal (Saphire lab, custom made VP40-2, 1:500) were used as primary antibodies. Secondary antibodies included Alexa 488-anti-mouse IgG (TFS, #A32723, 1:1000), Alexa 488-anti-rabbit IgG (TFS, #A11008, 1:1000), Alexa 647-anti-rabbit IgG (TFS, #A32733 1:1000), Alexa 568 anti-mouse IgG (TFS, #A11031, 1:1000). Nuclei were stained with Hoechst 33342 (TFS, #H3570) according to the manufacturer’s protocol. Coverslips were mounted with ProLong™ Gold Antifade Mountant (Invitrogen #P36930). Wide-field microscopy was performed with a KEYENCE BZX Microscope with a PlanApo 20 X objective lens (NA 0.75) and Plan Apo 100 X Oil objective lens (NA 1.45). Images were viewed, analyzed, and prepared for figures using Fiji software^61^.

#### Ultra-thin resin-embedded transmission electron microscopy

At 72 hours post-transfection (HEK 293T cells transiently expressing viral proteins), or 24 hr after infection (EBOV-GFP-ΔVP30-infected cells), cells were fixed in 2.5% glutaraldehyde in 0.10 M sodium cacodylate buffer pH 7.4 (EMS #15950) for 5 minutes at room temperature and for at least 2 hours at 4 °C. Cells were scraped from the culture vessel and pelleted by centrifugation at 200 x g for 5 min. After washing with 0.10 M sodium cacodylate buffer pH 7.4 (SC buffer) + 20mM Glycine, the pellets were treated with 1% osmium in 0.10 M SC buffer for 1 hour on ice and washed in 0.10 M SC buffer. The samples were incubated in 2% uranyl acetate for 1 hour at 4 °C and sequentially dehydrated with a 50, 70, 90, and 100% ethanol series followed by acetone. Then the samples were incubated in 50:50 Ethanol: Durcupan (Sigma Aldrich #44610) for >1 h at room temperature and in 100% Durcupan overnight. Pellets embedded in Durcupan were placed in a mold and cured in a 60 °C oven for 48 h. Ultrathin sections (60 nm) of the resin-embedded pellets were cut using a Leica ultramicrotome with a diamond knife. The sections were post-stained with both uranyl acetate and lead citrate. Images were captured on a JEOL 1400 plus TEM at 80 kV with a Gatan 4k × 4k camera and analyzed and prepared for figures using IMOD^62^.

#### Grid preparation

HEK 293T cells were transfected as indicated above. At three days after transfection the cells were detached and counted with a hemocytometer. Quantifoil 200 mesh holey carbon R1/4 copper grids (Quantifoil Micro Tools) were glow-discharged for 15 s at 0.3 mbar with 15 mA using a PELCO easiGlow glow discharge system (Ted Pella). A poly-lysine solution (2 µl; Sigma #4707) was applied to the grids immediately before 2,000-6,000 cells were pipetted onto the grid. Grids were blotted manually from the non-cell-deposited side using #1 Whatman filter paper. Plunging was performed in a humidity-controlled room (35% relative humidity) using a custom-made plunger to plunge the grids into a 50/50 mixture of liquid ethane and propane (Airgas) cooled to liquid nitrogen temperature. For EBOV-GFP-ΔVP30-infected cells, VP30-Vero cells (0.7-2 x 10^5^ cells/well) were seeded onto 35 mm dish containing Quantifoil 200 mesh holey carbon R1/4 or R2/2 gold grids (Quantifoil Micro Tools) one day before infection. EBOV-GFP-ΔVP30 infection was performed as indicated above. At 24 hours post-infection, cells were fixed with 4% PFA in 0.10 M sodium cacodylate buffer pH 7.4 for 15 min and plunged as described above. Just before plunging BSA Gold Tracer EM-grade 10 nm (EMS #25487) suspended in cell culture media was applied to the grids. The grids were clipped onto Cryo-FIB Autogrids (TFS). The samples were kept at liquid nitrogen temperature throughout the experiments.

#### Cryo-fluorescence microscopy

For cryo-fluorescence microscopy of EBOV-GFP-ΔVP30-infected cells, clipped grids were observed with a Leica EM Cryo CLEM system (Leica) using an HCX PL APO 50X objective lens with a numeric aperture of 0.90 cryo CLEM. Data was acquired using a matrix screener according to the manufacturer’s protocol and exported to MAPS format to open with Maps software (TFS) in the FIB-SEM microscope and TEM.

#### Cryo-focused ion beam milling

Cryogenic-focused ion beam milling was performed as described^27^ either manually using an Aquilos cryo-FIB–SEM microscope (TFS) or automatically using an Aquilos II cryo-FIB–SEM microscope (TFS). The locations of viral replication factories identified as GFP-positive regions outside the nucleus in EBOV-GFP-ΔVP30-infected cells were identified through correlation with SEM images obtained in Aquilos and a montage map generated using Leica Cryo-CLEM using global alignment function in Maps software. This approach allowed targeted milling of lamellae, including viral replication factories. For the transfected cells, we created the lamella randomly, and viral replication factory-like structures were visually identified during transmission electron microscopy observation. The thickness of lamellae used in this study was between 100 nm and 180 nm. Grids were stored in liquid nitrogen before imaging.

#### Cryo-electron tomography data acquisition

Tilt series were collected on a Titan Krios microscope (TFS) operating at 300 kV using a BioQuantum energy filter (slit width 20 eV) and a K3 direct electron detector (Gatan). The grids were loaded on the cassettes to align the lamella milling direction perpendicular to the tilting axis as described^27^. Maps software was used to acquire low-magnification montage maps for entire grids, and middle-magnification montage maps for individual lamella or cell periphery to visualize emerging virus particles with high defocus (−60 to −80 µm defocus). Correlation of the fluorescence images of the infected cells obtained by Leica cryo-CLEM was done using the global alignment function in Maps software. SerialEM^63^ was used to acquire tilt series data in low-dose mode. The parameters are summarized in Table S2.

#### Tomogram reconstruction and nucleocapsid annotation

Projection movie frames were imported to Warp software^64^ (v1.0.9) for gain reference and beam-induced motion correction, as well as contrast transfer function (CTF) and astigmatism estimation. Before tomogram reconstruction in Warp, per-tilt averaged images were imported to Etomo in IMOD package (v 4.11) for tilt series alignment. Alignment of tilt-series images was performed with patch-tracking, even for the cell periphery of non-milled samples, since the density of gold fiducials was insufficient for tracking. Tilt series that were poorly aligned and bad tilt images were manually removed. Reconstruction was done in Warp by importing the transformation files from Etomo. Deconvolved tomograms were also reconstructed in Warp. We determined the coordinates of the assembled nucleocapsids using fiber tracing modules in Amira 2021.1 (TFS). Before importing the 4-binned deconvolved tomograms into Amira, the file size was reduced using the IMOD newstack command. The approximate outer and inner diameters of intracellular nucleocapsids were measured from raw tomograms and used as input for the tracing parameters. After semi-automatic fiber tracing, nucleocapsid annotations in the tomograms were visually inspected and manually curated using the Amira filament editor. The resulting center coordinates were re-sampled equidistantly at every 4 nm using a script written in MATLAB^65,66^. For each point along the filaments, initial Euler angles were assigned for subsequent subtomogram analysis (see below).

#### Subtomogram analysis

Based on the resampled coordinates, we extracted subtomograms using 4-binned non-deconvolved tomograms using the Dynamo^67^ cropping function. The Euler angle orthogonal to the axis of the filament, i.e., the angle around the nucleocapsid, was randomized to minimize the effect of the missing wedge during cropping. An initial reference was generated by averaging subtomograms without alignment. A cylindrical mask was generated in Dynamo according to the initial average and used as an alignment mask. Other masks were left as defaults. Three iterations of translational and orientational alignment of the first two Euler angles, followed by six iterations of translational alignment and orientational alignment, including the third Euler angle around the axis of the nucleocapsid, with the third angle restricted to 20°, were performed. Since nucleocapsids are polar structures, cone-flip options in the Dynamo alignment parameter were selected during alignment. We estimated the helical parameters of the final Dynamo average using HI3D^68^. From the Dynamo-refined table, we removed junk and duplicated particles with a cross-correlation score followed by distance selection in Dynamo^69^. We calculated new coordinates and initial Euler angles based on the helical parameters along the helical axis centering nucleocapsid and created a new dynamo table using a script written in MATLAB^66^. We converted the dynamo tables to RELION files using a script, *dynamo2warp,* found in the dynamo2m package^70^. We extracted 4-binned subtomograms centering the nucleocapsid double layers in the center of the box (box size 42-43 nm) in Warp. We created initial templates using the *relion_reconstruct* function and performed 3D refinement using RELION v3.1^71^. The resulting density map was used as a reference for 3D classification without alignment into three classes. Class averages were inspected in UCSF ChimeraX^72^ and IMOD. Among the three classes, two classes lacked part of the outer nucleocapsid layers, when visualized with a higher threshold as compared to the selected class, as shown in Figures S2E and S2P. The particles from the selected class were re-extracted as 2-binned subtomograms in Warp. We first performed 3D refinement with RELION with a 240 Å diameter circular mask, followed by additional 3D refinement with a soft mask covering the nucleocapsid. We repeated this procedure with unbinned data. The resulting coordinates of each particle were then imported into M^73^ to perform iterative reference-based tilt-series refinement to generate high-resolution maps of the intracellular nucleocapsid. Gold-standard Fourier Shell Correlation plots were obtained with the RELION Post-processing module. Local resolution estimation and sharpening were done in M, and maps were rendered with ChimeraX.

For the intracellular nucleocapsid-like structure of NPΔ601-739-VP24-VP35, we repeated several rounds of RELION 3D refinement/M refinement, with recentering of the region of interest to obtain the maps having the best resolution of the outer nucleocapsid layer.

For the intracellular nucleocapsid-like structure of NP-VP24-VP35, resolution achieved with 3D refinement of unbinned data was not higher than that with 2-binned data. As such, we performed refinement steps with M as 2-binned data. After M, we repeated one round of RELION 3D refinement.

#### Heterogeneous population analysis of intracellular nucleocapsids and visualization of particle distribution

We obtained an intracellular, fully assembled nucleocapsid structure at 9 Å resolution from cells expressing NPΔ601-739, VP24, and VP35 (Figure 2). To reveal the heterogeneity of intracellular nucleocapsids, we gathered particles sorted as irregular nucleocapsid classes during the RELION 3D classification (Classes 1 and 2 in Figure S2E) and re-classified into 10 classes by 3D classification without alignment (Now Classes 1-10 in Figure S2W). The position of the individual particles are plotted back to the one representative tomogram using ArtiaX^74^.

#### Measuring distances/angles between intracellular nucleocapsid

Measuring the angles and distances between intracellular nucleocapsids was done as described^66^. Briefly, we measured distances between points sampled every 4 nm along each nucleocapsid and other nucleocapsids (see Tomogram reconstruction and nucleocapsid annotation). We focused on point pairs separated by less than 100 nm. For each point, we computed (1) a tangent vector along the nucleocapsid, (2) the distance to the nearest point in neighboring filaments, and (3) the angle between these vectors (0° indicating parallel). We calculated and visualized the data in a 2-D histogram using MATLAB.

#### Mapping the average back into 3D volume

Mapping the average back to the tomogram volume shown in Figure 3 and Video S1-2 was done with the *dynamo_table_chimera* function using dynamo tables and averages obtained after Dynamo averaging. The map was visualized with Chimera^75^ and movies were created with Amira.

#### Tomogram segmentation

For visualization of densities in viral replication factories, fully assembled intracellular nucleocapsids and membranes in the tomogram shown in Figures 5E, threshold segmentation using magic wand tool, fiber tracing modules in Amira and MemBrain^76^ were used for annotation respectively. Each output was inspected, and cleaned manually, and the final figure was generated with Amira.

#### Model building

Many structures were available for individual Ebola nucleocapsid components. We started modeling the intracellular nucleocapsid-like structure of NPΔ601-739-VP24-VP35 using a local sharpened map obtained after M processing. The map was further modified in OccuPy^77^ to equalize the map according to occupancy to facilitate model fitting into the map based on the shape rather than density gradient. The following structures were used for rigid body fitting using ChimeraX: in-virion Ebola virus nucleocapsid model composing two NP cores and two VP24 molecules (PDB: 6EHM)^25^, NP core (PDB: 6EHL)^25^, NP bound to VP35 chaperone peptide (PDB: 4ZTG^11^, 4YPI^12^), VP24 (PDB: 4M0Q^30^), VP35 C-terminal domain (PDB: 3FKE^31^) and the VP35 protein and polymerase L from the EBOV L-VP35 complex (PDB: 7YER^15^, 8JSL, 8JSM, 8JSN^51^). Based on the in-virion Ebola virus nucleocapsid model comprising 2NP and 2VP24 (PDB: 6EHM), locations of two NP core structures (6EHL) and two VP24 (4M0Q) were located in the central repeating unit of the EM map using ChimeraX *Fit in Map* function. The remaining unfitted density was sequentially fitted with all protein components of the EBOV virion, either fragments or full sequences from the above mentioned PDBs. The N-lobe of the NP core (6EHL) and VP35 C-terminal domain (3FKE) fitted well in the larger and smaller densities, respectively, that were attached to two VP24 molecules. Visual analysis of these fits showed fitting of secondary structure elements not only in the central repeating unit but also in adjacent repeating units in the map, confirming the fit.

For outer NP, we compared fitting of the NP-core structure with the other two structures of NP bound to the VP35 N-terminal NP binding peptide (4ZTG and 4YPI). Both structures fit better to the map than the initially placed NP core (6EHL) since the density corresponding to VP35 binding peptides was visible in our EM map but missing in the NP core structure (6EHL), as expected. Comparison with 4ZTG and 4YPI showed that the part of the long alpha helix (aa 355-385) in the C-lobe of 4YPI protruded from our map. This arrangement arises because 4YPI is part of the oligomerized structure^12^, and oligomerization might stabilize this helix similar to NP forming NP-RNA central core^25^. In contrast, 4ZTG, which behaves as a monomer in solution and the corresponding part of the helix was unstable, therefore was not modeled^11^. Considering that outer NP is monomeric, we chose 4ZTG for further flexible fitting. We found that the assembled model also fit with our intracellular nucleocapsid-like structure of full-length NP and in-virion nucleocapsid structure from virus-like particles (EMD-3781) and Ebola virus (EMD-3873)^25^. In this study, for NP-core structures (PDB: 6EHL) and outer NP bound with VP35 peptide (PDB: 4ZTG), we used AlphaFold 2^78^, integrated within the ColabFold notebook, to generate protein models in the PDB format. This approach was necessary due to the incompatibility of available online PDB structures with the force field used for the flexible refine strategy. We used PDB ID as a single template and we confirmed the general geometry of the prediction with the template. This comparison verified that the AlphaFold 2 models were structurally congruent with the original PDB models. Next, we used NAMDinator^79^, an automated molecular dynamics flexible fitting pipeline, to create two refined models, one for our intracellular nucleocapsid EM map derived from NPΔ601-739 and the other for in-virion nucleocapsid-like structure (EMD-3781)^25^. The model assembly was performed with flatten_obj in PyMol (Schrödinger). We used Phenix Refine 1.13^80^ at the end of NAMD flexible fitting for refinement steps and obtained two individually refined models, the intracellular nucleocapsid and the in-virion nucleocapsid models. Due to map resolution, we truncated models to poly-glycine chains using PDB tools in the Phenix package to avoid overfitting the model. We noted there is an unassigned density indicated with black arrows in Figures S3I and S3J. This density might correspond to the part of the unstructured C-terminal region of VP24 (aa 232-251), which has not been modeled in pre-existing VP24 crystal structure PDB: 4M0Q^30^ used for our integrative modeling. The VP24 (aa 231) modeled in our molecular model, locates close to this density. However, due to the lack of sufficient resolution with our current EM map, we decided not to model this identity. The final statistics reported in Table S3 are reported for the poly-glycine model.

By fitting the refined intracellular nucleocapsid model into the several repeating units of intracellular nucleocapsid structure obtained from NPΔ601-739, we estimated the region of N-terminal parts of VP35. For that, we fit the longest modeled VP35 protein structure (aa 81 to 340) of the polymerase L-VP35 complex (7YER 8JSL, 8JSM, 8JSN)^15,51^ into region corresponding to VP35 CTD using the *MatchMaker* function of ChimeraX. These VP35 models showed some variation in the angle of the oligomerization domain, but the N-terminal region (aa 81 to 110) from all these models pointed through the outer NP density of the adjacent repeating unit. We chose 7YER as a representative structure to model hypothetical N-terminal parts of VP35 fitted in the nucleocapsid structure.

#### Interface analysis

Interfaces were identified by visually examining the atomic model fitted in the experimental map using the following criteria: (1) densities corresponding to the specified loops or helices are visible in the EM map; and (2), even increasing the threshold of the density, the connecting density between two proteins is still visible, assuming the regions forming interfaces are more rigid than elsewhere. We did not consider a region as an interface if the corresponding experimental EM densities were not observed, even if the distance shown in the model was small. We also observed potential interfaces where experimental density showed connections, but no specific residues were assigned in the atomic model.

#### Sequence conservation analysis

Protein sequences of NP, VP24, and VP35 from six filoviruses, Zaire Ebola virus, Bundibugyo virus, Tai Forest virus, Sudan virus, Marburg virus, and Lloviu virus were retrieved from the National Center for Biotechnology Information (NCBI) GenBank, and multiple sequence alignment was performed with the T-Coffee package^81^. We calculated a conservation score for each position using three different metrics, Livingstone, Karlin, and Valdar^41–45^, which have different properties. We calculated all non normalized conservation scores by running AACON from jalview Web service (https://www.compbio.dundee.ac.uk/aacon/) by keeping all parameters default. Briefly, the Livingstone metric is purely based on the conservation of physicochemical properties, whereas the Karlin metric is based on an evolutionary substitution matrix, and the Valdar metric is a hybrid of the two. The maximum conservation score, assigned when a given amino acid is identical in all 6 filoviruses considered, is 1 and 11 for the Karlin and Livingstone metrics, respectively. The maximum value for the Valdar conservation score ranged between 8 to 13. We compared the average conservation score of each interface region with that from full-length proteins.

### QUANTIFICATION AND STATISTICAL ANALYSIS

The details of the quantification and all statistical analyses are included in figure legends or the relevant section of methods details.

## Supplemental videos and tables

**Video S1.** *In situ* cryo-FIB-ET tomogram of HEK 293T cell expressing full-length NP, VP24 and VP35 and mapping back the average of nucleocapsid-like structure to the tomogram, related to Figures 1 and 3.

**Video S2.** *In situ* cryo-FIB-ET tomogram of HEK 293T cell expressing NPΔ601-739, VP24 and VP35 and mapping back the average of nucleocapsid-like structure to the tomogram, related to Figures 2 and 3.

**Video S3.** EM map of intracellular nucleocapsid-like structure revealed in cells expressing NPΔ601-739, VP24 and VP35 and structural model of intracellular nucleocapsid, related to Figures 2 and 6.

**Video S4.** Structural models showing intracellular nucleocapsid is further condensed upon incorporation into virion, related to Figures 4 and 7.

**Table S1.** Interaction sites involved in nucleocapsid assembly and average conservation scores obtained with three different scoring metrics, related to Figure 6

**Table S2.** Cryo-ET data collection and processing, related to Figures 1, 2, 3, and 5

**Table S3.** Refinement statistics, related to Figures 2, and 4

