## Supplementary material for "Intracellular Ebola Virus nucleocapsid assembly revealed by *in situ* cryo-electron tomography": Supple_figures and table

**Figure S1. NP and NPΔ601-739 colocalize with VP24 and VP35 and they form nucleocapsid-like structures in cells indistinguishable from EBOV-GFP-ΔVP30 virus-infected cells**

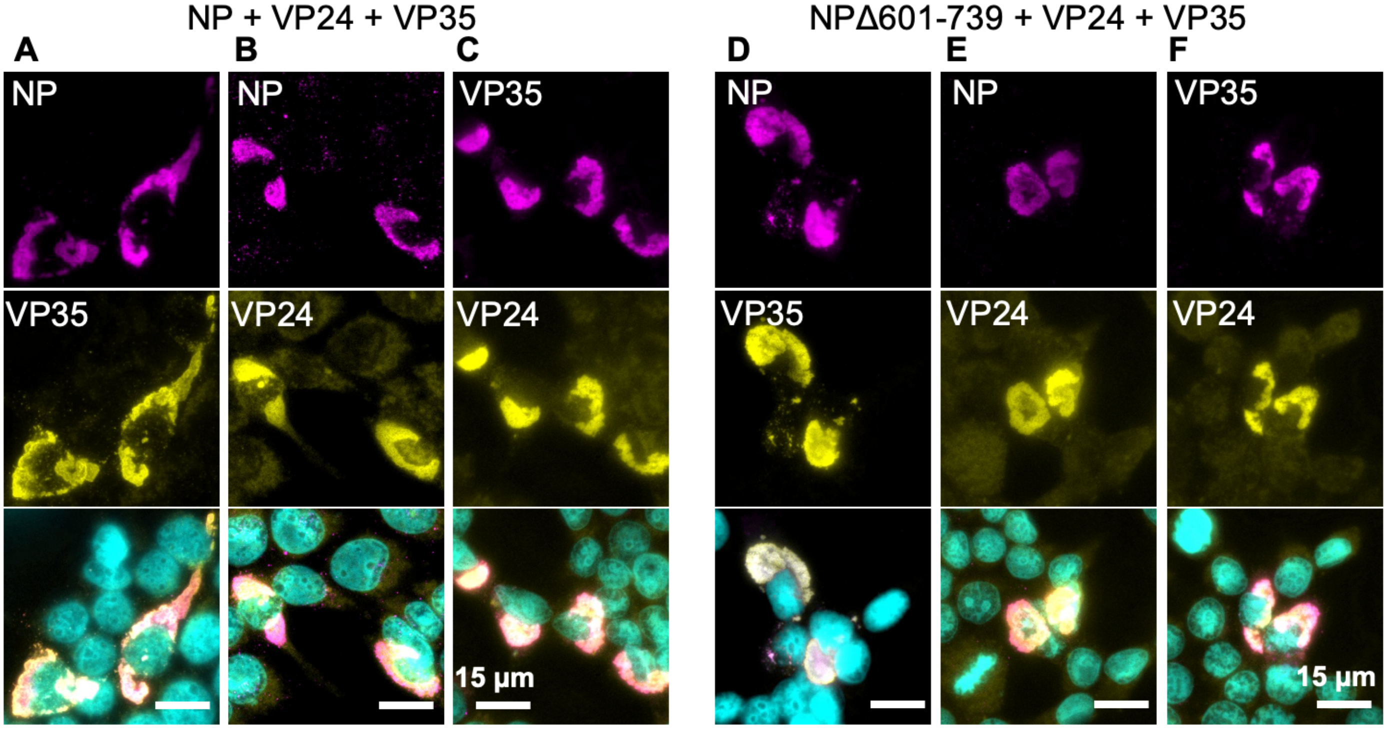

**Figure S1 (continue). NP and NPΔ601-739 colocalize with VP24 and VP35 and they form nucleocapsid-like structures in cells indistinguishable from EBOV-GFP-ΔVP30 virus-infected cells**

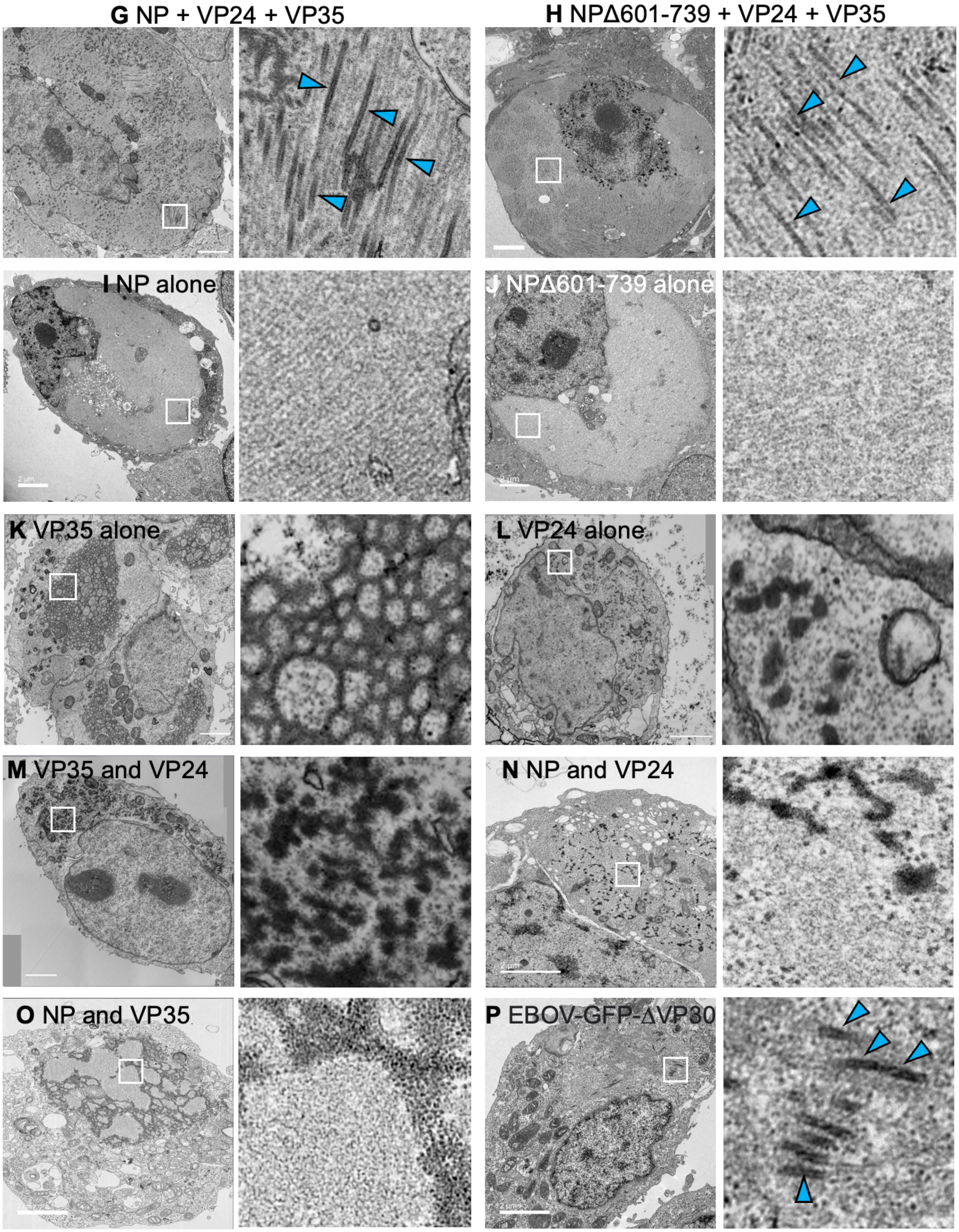

**Figure S2. Subtomogram averaging**

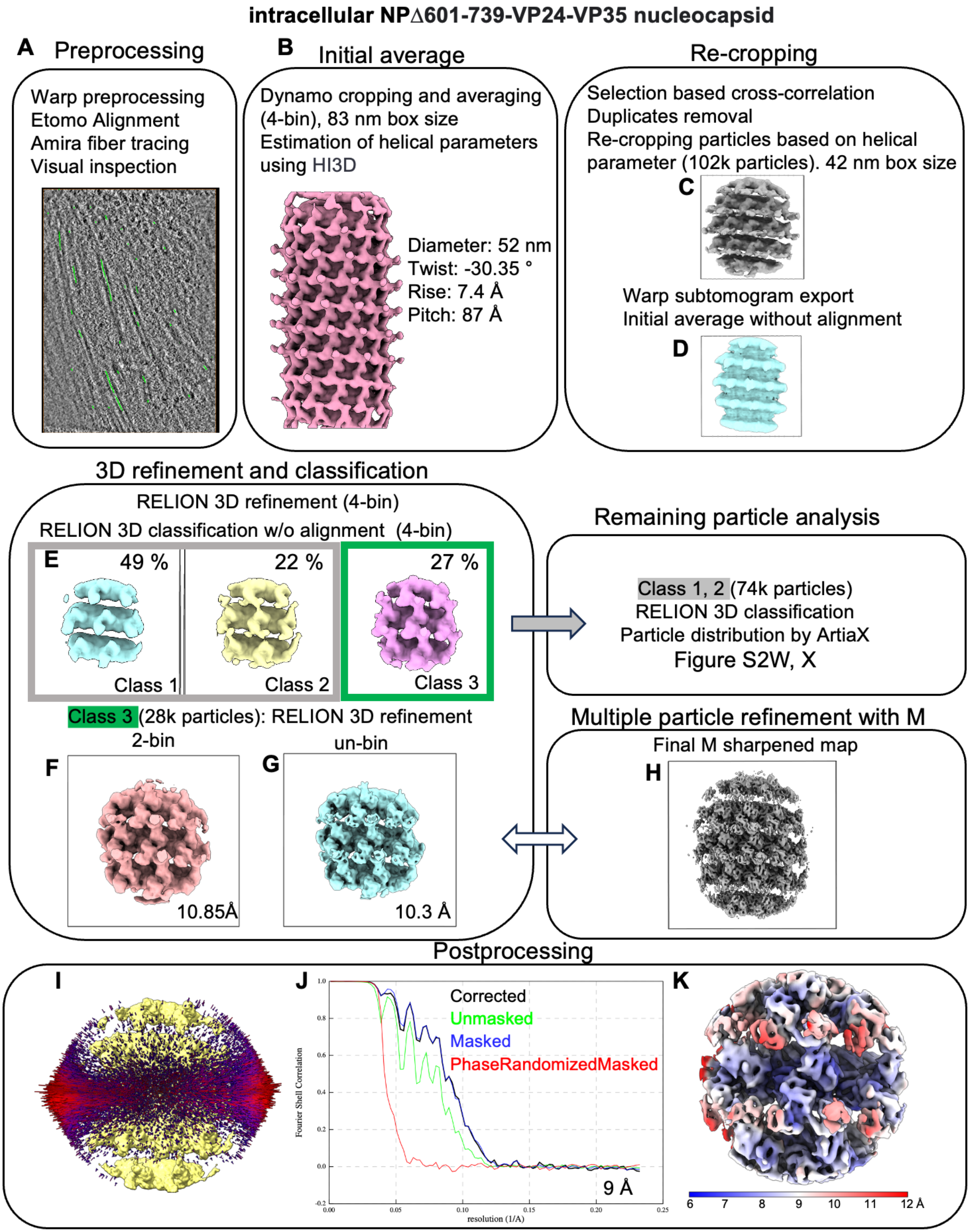

**Figure S2(continue). Subtomogram averaging**

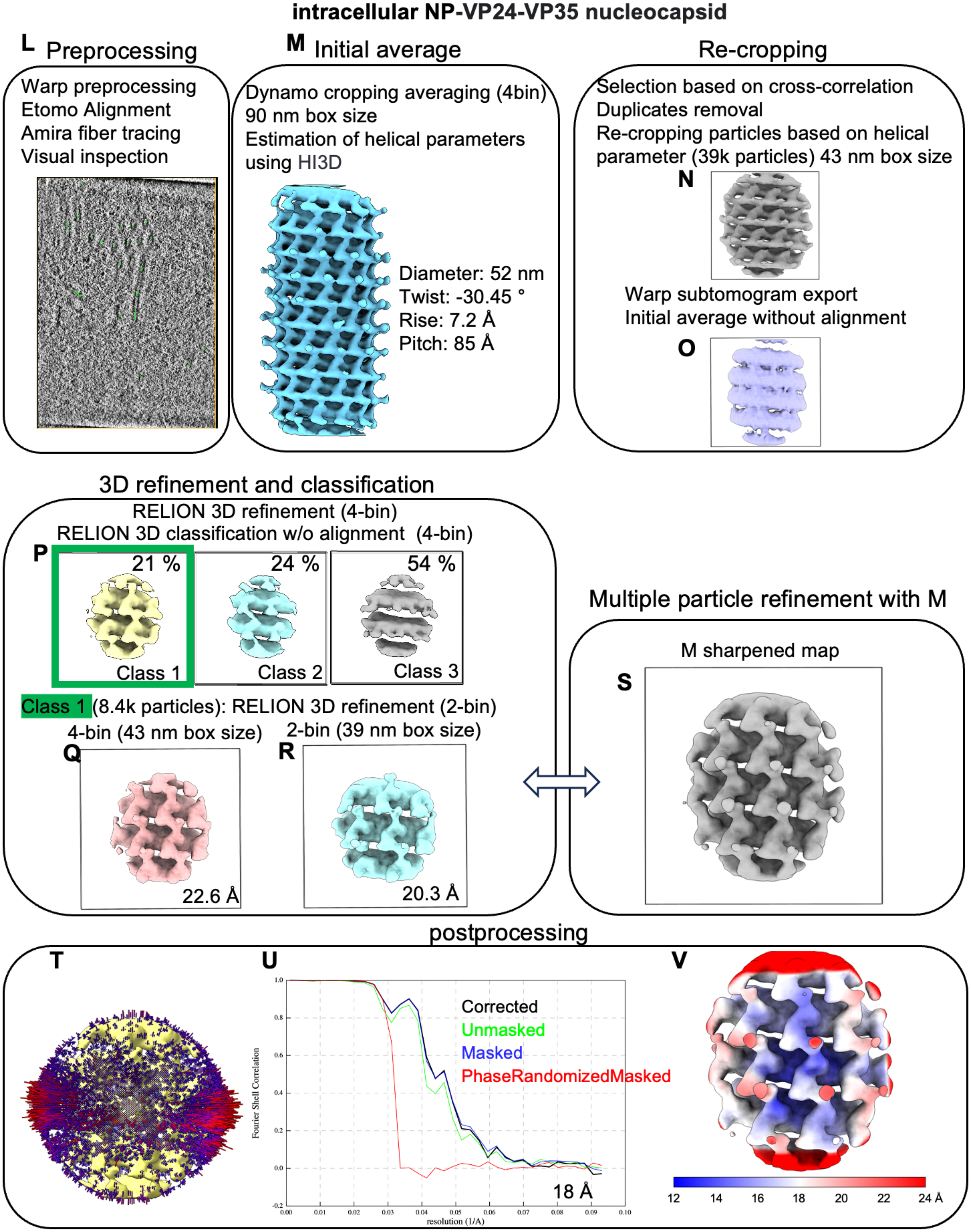

**Figure S2(continue). Subtomogram averaging**

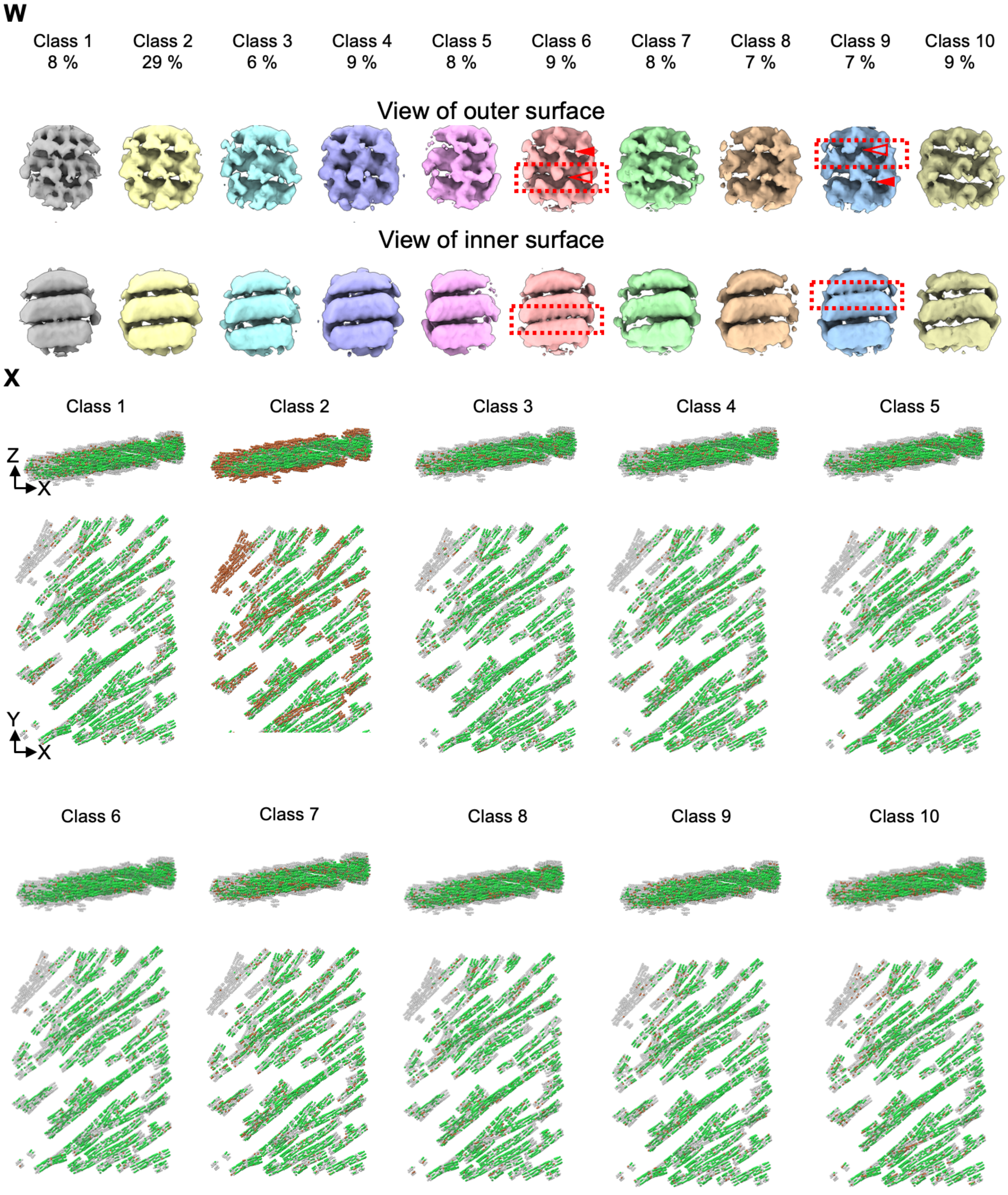

**Figure S3. Model building of intracellular Ebola virus nucleocapsid**

**A.** Structure used for integrative modeling

| Model | Protein | PDB entry | Modeled region indicated by residue number |
|---|---|---|---|
| | 2NP_2VP24 | 6EHM | NP; 16-404, VP24; 11-231 |
| | NP | 6EHL | 16-405 |
| | NP complex with VP35 N-terminal binding peptide | 4ZTG | VP35;22-45 , NP; 36-354 |
| | NP complex with VP35 N-terminal binding peptide | 4YPI | VP35;20-47 , NP; 38-385 |
| | VP24 | 4M0Q | 11-231 |
| | VP35 C-terminal domain | 3FKE | 218-340 |
| | VP35 complex with polymease L | 7YER | VP35; 81-340, polymerase L; 8-1383 |

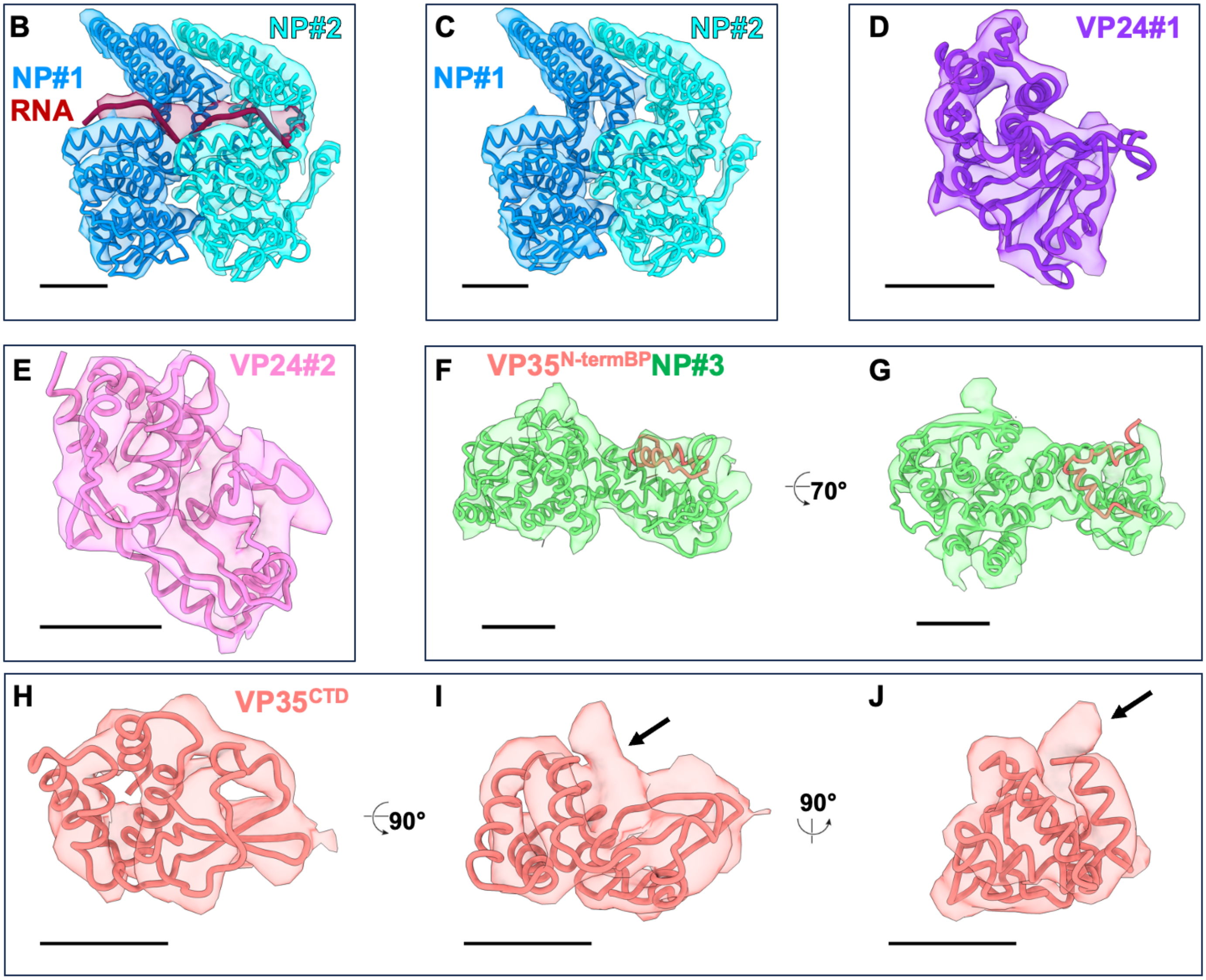

**Figure S4. Comparison of various nucleocapsid structures**

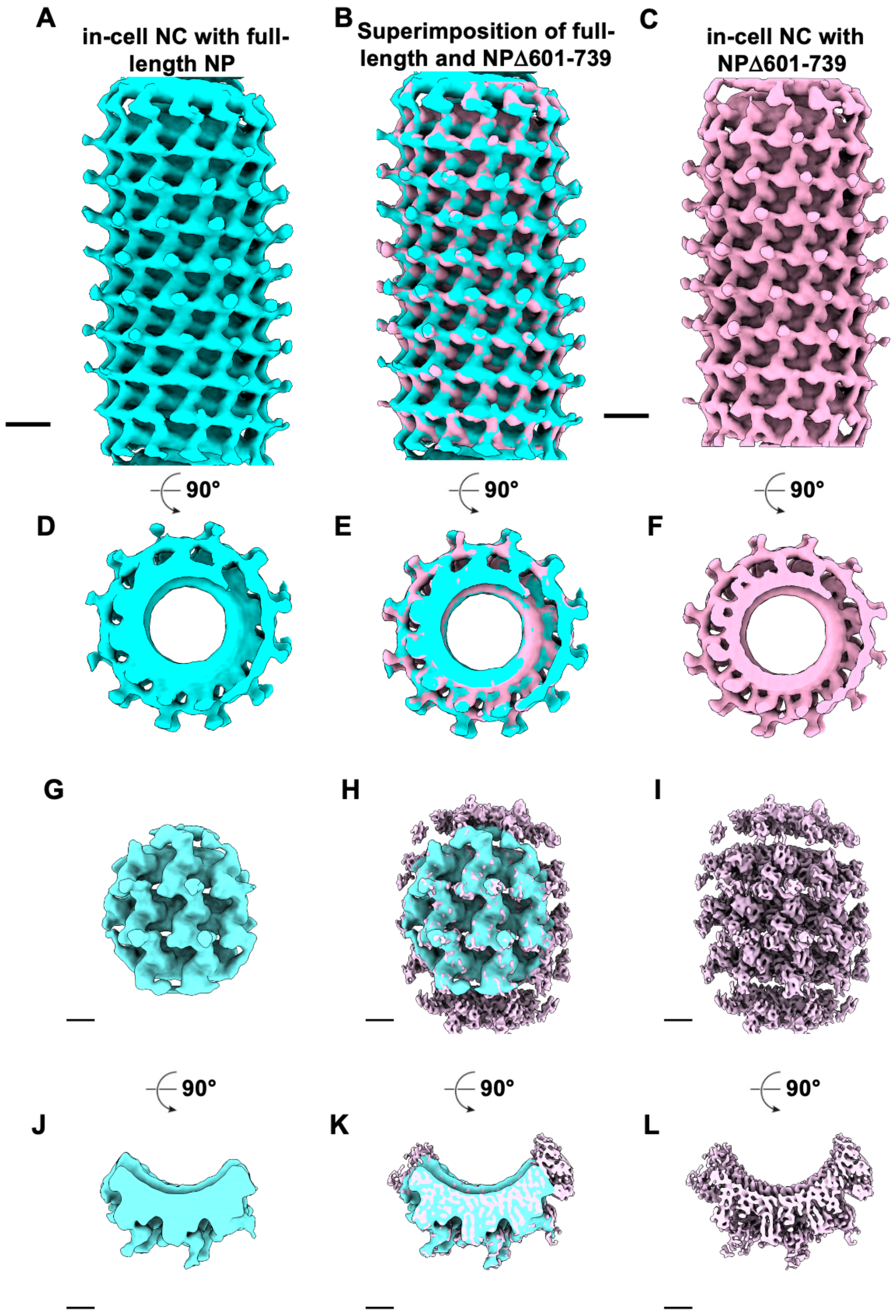

**Figure S4 (continue). Comparison of various nucleocapsid structures**

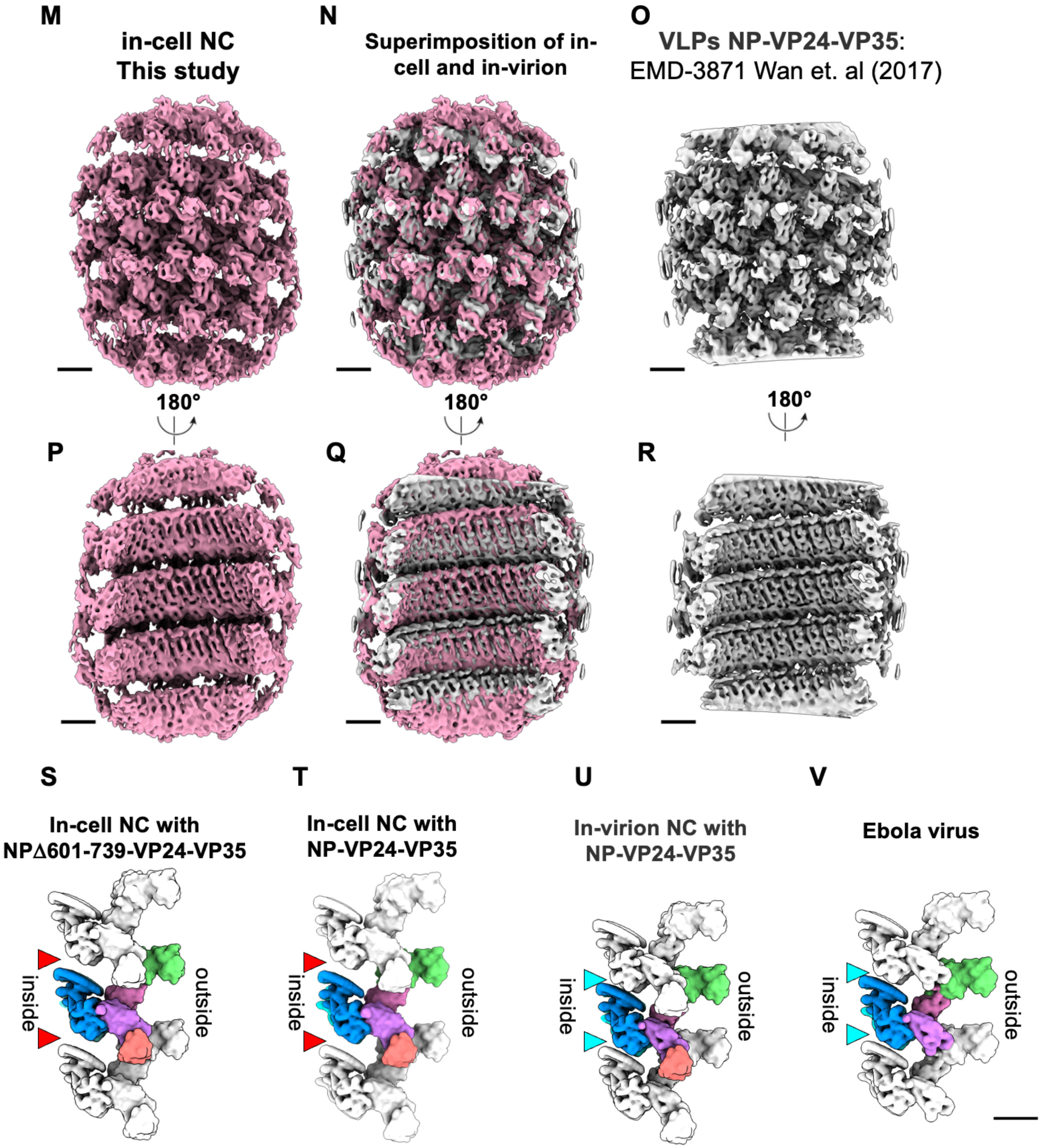

**Figure S5. Visualization of EBOV-GFP-ΔVP30-infected cells**

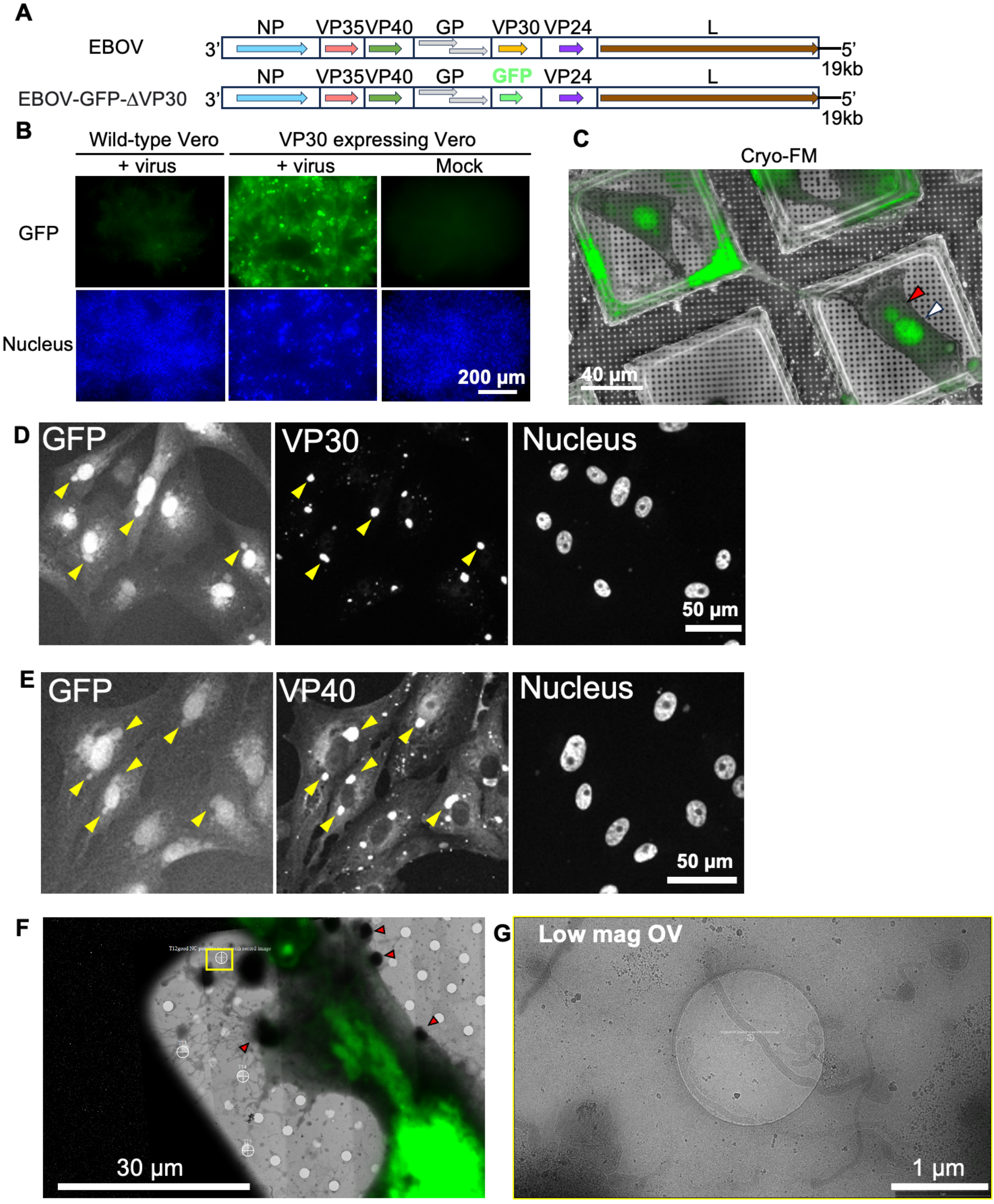

**Figure S6. Sequence alignment of orthologs of NP, VP24, and VP35 and regions involved in the interface and average conservation scores**

**A** NP

```
AI011747.1      MNSRPQKVWMTPSLTESDMDYHKILTAGLSVQQGIVRQRVIPVYQVNNLEEICQLIIQAFEAGVDFQESADSFLLMLCLH
QYU59244.1      MDPRPIRTWMMHNTSEVEADYHKILTAGLSVQQGIVRQRIIPVYQISNLEEVCQLIIQAFEAGVDFQDSADSFLLMLCLH
ACI28629.1      MESRAHKAWMTHTASGFETDYHKILTAGLSVQQGIVRQRVIQVHQVTNLEEICQLIIQAFEAGVDFQESADSFLLMLCLH
QDC12555.1      MDKRVRGSWALGGQSEVDLDYHKILTAGLSVQQGIVRQRVIPVYVVNDLEGICQHIIQAFEAGVDFQDNADSFLLLLCLH
AJD39282.1      ------------------MDLHSLLELGTKPTAPHVRNKKVILFDTNHQVSICNQIIDAINSGIDLGDLLEGGLLTLCVE
UKR35326.1      MNRYLGHGTRTSRENTNLSELHGILSLGLNVDHTIVRKKSIPLFEIGNSDQVCNWIIQIIEAGVDLQDVADSFLTMLCVN
                                   : * :*  * .     **:: : :.   .   :*: **: :::*:*: :  :. *  **:.

AI011747.1      HAYQGDYKLFLESGAVKYLEGHGFRFEVKKRDGVKRLEELLPAVSSGKNIKRTLAAMPEEETTEANAGQFLSFASLFLPK
QYU59244.1      HAYQGDYKQFLESNAVKYLEGHGFRFEMKKKEGVKRLEELLPAASSGKNIKRTLAAMPEEETTEANAGQFLSFASLFLPK
ACI28629.1      HAYQGDYKQFLESNAVKYLEGHGFRFEVRKKEGVKRLEELLPAASSGKSIRRTLAAMPEEETTEANAGQFLSFASLFLPK
QDC12555.1      HAYQGDHRLFLKSDAVQYLEGHGFRFEVREKENVHRLDELLPNVTGGKNLRRTLAAMPEEETTEANAGQFLSFASLFLPK
AJD39282.1      HYYNSDKDKFNTSPIAKYLRDAGYEFDVIKNADATRFLDVIPNEPHYSPLILALKTLESTESQRGRIGLFLSFCSLFLPK
UKR35326.1      HAYQGDPNLFLESPAAHYLKGHGIHFEIQHRDNVDHITDLLGVGSRDKSLRKTLSALEFEPDGSTTAGMFLSFASLFLPK
                * *:.*   *  *  .:**.. * .*:: .. .. :: :::   .  . :  :* ::          * ****.******

AI011747.1      LVVGEKACLEKVQRQIQVHAEQGLIQYPTAWQSVGHMMVIFRLMRTNFLIKFLLIHQGMHMVAGHDANDAVISNSVAQAR
QYU59244.1      LVVGEKACLEKVQRQIQVHAEQGLIQYPTSWQSVGHMMVIFRLMRTNFLIKFLLIHQGMHMVAGHDANDAVIANSVAQAR
ACI28629.1      LVVGEKACLEKVQRQIQVHSEQGLIQYPTAWQSVGHMMVIFRLMRTNFLIKFLLIHQGMHMVAGHDANDAVIANSVAQAR
QDC12555.1      LVVGEKACLEKVQRQIQVHAEQGLIQYPTSWQSVGHMMVIFRLMRTNFLIKFLLIHQGMHMVAGHDANDTVISNSVAQAR
AJD39282.1      LVVGDRASIEKALRQVTVHQEQGIVTYPNHWLTTGHMKVIFGILRSSFILKFVLIHQGVNLVTGHDAYDSIISNSVGQTR
UKR35326.1      LVVGERACLEKVQRQIQIHAEQGLIQYPTQWQSVGHMMVVFRLIRVNFVLKFLLVHQGMHMMAGHDANDAIIANSISQTR
                ****::*.:**. **: :* ***:: **. * :.*** *:* ::* .*::**:*:***::::**** *::*:**:.*:*

AI011747.1      FSGLLIVKTVLDHILQKTERGVRLHPLARTAKVKNEVNSFKAALSSLAKHGEYAPFARLLNLSGVNNLEHGLFPQLSAIA
QYU59244.1      FSGLLIVKTVLDHILQKTEHGVRLHPLARTAKVKNEVSSFKAALASLAQHGEYAPFARLLNLSGVNNLEHGLFPQLSAIA
ACI28629.1      FSGLLIVKTVLDHILQKTEHGVRLHPLARTAKVKNEVNSFKAALSSLAQHGEYAPFARLLNLSGVNNLEHGLFPQLSAIA
QDC12555.1      FSGLLIVKTVLDHILQKTDLGVRLHPLARTAKVKNEVSSFKAALGSLAKHGEYAPFARLLNLSGVNNLEHGLYPQLSAIA
AJD39282.1      FSGLLIVKTVLEFILQKTDSGVTLHPLVRTSKVKNEVASFKQALSNLARHGEYAPFARVLNLSGINNLEHGLYPQLSAIA
UKR35326.1      FSGLLIVKTVLEHILQKTEAGVQLHPLARTSKVKGELLAFKSALEALASHREYAPFARLLNLSGVNNLEHGLYPQLSAIA
                ***********:.*****: ** ****.**:***.*: :** **  ** * *******:*****:*******:*******

AI011747.1      LGVATAHGSTLAGVNVGEQYQQLREAATEAEKQLQQYAESRELDHLGLDDQEKKILMNFHQKKNEISFQQTNAMVTLRKE
QYU59244.1      LGVATAHGSTLAGVNVGEQYQQLREAATEAEKQLQKYAESRELDHLGLDDQEKKILKDFHQKKNEISFQQTTAMVTLRKE
ACI28629.1      LGVATAHGSTLAGVNVGEQYQQLREAATEAEKQLQKYAESRELDHLGLDDQEKKILKDFHQKKNEISFQQTTAMVTLRKE
QDC12555.1      LGVATAHGSTLAGVNVGEQYQQLREAATEAEKQLQQYAETRELDNLGLDEQEKKILMSFHQKKNEISFQQTNAMVTLRKE
AJD39282.1      LGVATAHGSTLAGVNVGEQYQQLREAAHDAEVKLQRRHEHQEIQAIAEDDEERKILEQFHLQKTEITHSQTLAVLSQKRE
UKR35326.1      LGVATAHGSTLAGVNVSEQYQQLREAATEAEKQLQQHSEMRELETLGLDEQERKILATFHSRKNEINIQQTSSILAIRKE
                ****************.********** :** :**:  * :*:: :. *::*:***  ** :*.**. .** :::: ::*

AI011747.1      RLAKLTEAITAASLPK---TSGHYDDDDDIP-----FPGPINDDDNPGHQDDDPTDSQDTTIPDVVVDPDDGSYGEYQSY
QYU59244.1      RLAKLTEAITSTSILK---TGRRYDDDNDIP-----FPGPINDNENSGQNDDDPTDSQDTTIPDVIIDPNDGGYNNYSDY
ACI28629.1      RLAKLTEAITSTSLLK---TGKQYDDDNDIP-----FPGPINDNENSEQQDDDPTDSQDTTIPDIIVDPDDGRYNNYGDY
QDC12555.1      RLAKLTEAITTASKIK---VGDRYPDDNDIP-----FPGPIYDDTHPNPSDDNPDDSRDTTIPGGVVDPYDDESNNYPDY
AJD39282.1      KLARLAAEIENNIVED---QGFKQS-QNRVSQSFLNDPTPVEVTVQ------ARPVNRPTALPPPVDDKIEHESTGDSSS
UKR35326.1      RLRKLTEALTEEKNKNALDDEDESEEDDWSP-----ENRGIRSNKGSTKEPSSYTASRTEEDRNNYNSKDD-HLSGKEQM
                :* :*:  :      .      .   ::  .         :               .:         .  :       .

AI011747.1      SENGMNAPDDLVLFDLDEDDED-TKPVPNRSTKGGQQKNSQ--KGQHTEGRQTQS-RPTQNVPGPHR-----AIHHTSAP
QYU59244.1      ANDAASAPDDLVLFDLEDEDDA-DNPAQNTPEKNDRPATTKLRNGQDQDGNQGET-ASPRVAPNQ-------YRDKPMPQ
ACI28629.1      PSETANAPEDLVLFDLEDGDED-DHRPSSSSENNNKHSLT----GTDSNKTSNWN-RNPTNMPKKDST---QNNDNPAQR
QDC12555.1      EDSAEGTTGDLDLFNLDDDDDD-SRPGPPDRGQ-NKERAART-YGLQDPTLDGAK-KVPELTPGSHQ-----PGNLHITK
AJD39282.1      SSSFVDLNDPFALLNEDEDTLDDSVMIPSTTSREFQGIPES---------------------PGQSQ-----NLDDSQGK
UKR35326.1      STQQESGADDLDLFDLDDDGDT-NSQDPNNRQK-QSDTQQ----TQESSDRSDYSRRPAYDWPPGDRPHTTQATDEHTDL
                  .  .    : *:: ::              .                             *           .

AI011747.1      LTDNDRRNE--P-SSSTSPRMLTPINEEADPLDDADDETSSLPPLESDDEEQDRDGTSN-RTPTVAPPAPVYRDHSEKKE
QYU59244.1      VQDRSENHD--Q-TLQTQSRVLTPISEEADPSDHNDGDNESIPPLESDDEGSTDTTAAE-TKPATAPPAPVYRSISVDDS
ACI28629.1      AQEYARDNI--Q-DTPTPHRALTPISEETGSNGHNEDDIDSIPPLESDEENNTETTITT-TKNTTAPPAPVYRSNSEKEP
QDC12555.1      SGSNTNQPQ--G-NMSSTLHSMTPIQEESEPDDQKDNDDESLTSLDSEGDEDGESISEE-NTPTVAPPAPVYKDTGVDTN
AJD39282.1      QEDESTNPI-K--KQFLRYQELPPVQEDDES-EYTTDSQESIDQPGS----------DN-EQGVDLPPPPLYAQEKRQDP
UKR35326.1      LNKDHRRNQVKPGRRGNDPRTLPLISFD-DNEGEILDDKSDLPAPDTHSDPDTEESEEEHPDEELLPPAPKYSTKTSEQE
                  .                : :. :. :        .. ..:    :                   **.* *     .

AI011747.1      -----LPQDEQQDQDHTQEARNQD-----S-DNTQPEHSFEEMYRHILRSQGPFDAVLYYHMMKDEPVVFSTSDGKEYTY
QYU59244.1      -----VPSENIPAQSNQTNNEDNV-----R-NNAQSEQSIAEMYQHILKTQGPFDAILYYHMMKEEPIIFSTSDGKEYTY
ACI28629.1      -----LPQEKSQKQPNQVSGSENT-----D-NKPHSEQSVEEMYRHILQTQGPFDAILYYYMMTEEPIVFSTSDGKEYVY
QDC12555.1      -----QQNGPSSTVDSQGSESEAL-----P-INSKKSSALEETYYHLLKTQGPFEAINYYHLMSDEPIAFSTESGKEYIF
AJD39282.1      -----IQHPAASSQDPFGSIGDVNGDILEPIRSPSSPSAPQEDTRAREAYELSPDFTNYEDNQQNWPQRVVTKKGRTFLY
UKR35326.1      PGDWKQPTSPLSTIPEEEGGHEAN-----N-DNS-ESDLIDQMYQHIFKTEGAYAAINYYYKTTGRPVTFTSNNNHDYTF
                                     :          ..       :        : .     *       *  . :...: : :

AI011747.1      PDSLEEEYPPWLTEKEAMNEENRFVTLDGQQFYWPVMNHKNKFMAILQHHQ
QYU59244.1      PDSLEDEYPPWLSEKEAMNEDNRFITMDGQQFYWPVMNHRNKFMAILQHHR
ACI28629.1      PDSLEGEHPPWLSEKEALNEDNRFITMDDQQFYWPVMNHRNKFMAILQHHK
QDC12555.1      PDSLEEAYPPWLSEKEALEKENRYLVIDGQQFFWPVMSLQDKFLAVLQH-D
AJD39282.1      PNDLLQTSPPESLVTALVEEYQNPVSAKELQADWPDMSFDERRHVAMN--L
UKR35326.1      PQDTEGLFPPWEGKENQKVAEILTNSLHETGQEWADMSAKERYLFLIN--N
                *:.     **                 .     *. *.  ::    ::
```

Interface (i)

Interface (ii)

Interface (iii)

Interface (iv)

Interface (v)

Interface (vi)

Interface (vii)

AIO11747.1 NP [Zaire ebolavirus]
QYU59244.1 NP [Bundibugyo ebolavirus]
ACI28629.1 NP [Tai Forest ebolavirus]
QDC12555.1 NP [Sudan ebolavirus]
AJD39282.1 NP [Orthomarburgvirus marburgense]
UKR35326.1 NP [Cuevavirus lloviuense]

**Figure S6 (continue). Sequence alignment of orthologs of NP, VP24, and VP35 and regions involved in the interface and average conservation scores**

Interface (i)
Interface (ii)
Interface (iii)
Interface (iv)
Interface (v)
Interface (vi)
Interface (vii)

## B VP24

```
AAM76037.1      MAKATGRYNLISPKKDLEKGVVLSDLCNFLVSQTIQGWKVYWAGIEFDVIHKGMALLHRLKTNDFAPAWSMTRNLFPHLF
ACI28626.1      MAKATGRYNLVSPKKDLERGLVLSDLCTFLVDQTIQGWRVTWVGIEFDIAQKGMALLHRLKTADFAPAWSMTRNLFPHLF
ACI28635.1      MAKATGRYNLISPKKDLEKGLVLNDLCTLSVAQTVQGWKVTWAGIEFDVTQKGMALLHRLKTSDFAPAWSMTRNLFPHLF
AGL50930.1      MAKATGRYNLVTPKRELEQGVVFSDLCNFLVTPTVQGWKVYWAGLEFDVNQKGITLLNRLKVNDFAPAWAMTRNLFPHLF
CAA78119.1      MAELSTRYNLPANVTENSINLDLNSTARWIKEPSVGGWTVKWGNFVFHIPNTGMTLLHHLKSNFVVPEWQQTRNLFSHLF
UKR35333.1      MARPSGRYNILQTDTLHENTVVKHDLCNFLVTTTITGWDVYWAGHLFHVPNKGIALLTRLKTSDFAPAWTLTKNLFPHLF
                **. : ***:        .  :   . .      :: ** * * .  *.: :.*::** :**   ..* *  *:***.***

AAM76037.1      QNPNSTIESPLWALRVILAAGIQDQLIDQSLIEPLAGALGLISDWLLTTNTNHFNMRTQRVKEQLSLKMLSLIRSNILKF
ACI28626.1      QNSNSTIESPLWALRVILAAGIQDQLIDQSLVEPLAGALSLVSDWLLTTNTNHFQMRTQHAKEQLSLKMLSLVRSNILKF
ACI28635.1      QNPNSTIESPLWALRVILAAGIQDQLIDQSLIEPLAGALGLIADWLLTTGTNHFQMRTQQAKEQLSLKMLSLVRSNILKF
AGL50930.1      KNQQSEVQTPIWALRVILAAGILDQLMDHSLIEPLSGALNLIADWLLTTSTNHFNMRTQRVKDQLSMRMLSLIRSNIINF
CAA78119.1      KNPKSTIIEPFLALRILLGVALKDQELQQSLIPGFRSIVHMLSEWLLLEVTSAIHISPNLLGIYLTSDMFKILMAGVKNF
UKR35333.1      HNPTSTLKDPVWGLRVILAAALYDQVMEQSLIKPLSETLTLLGDWLLTTETQKFNLRTPRAQEQISLRMLSLVKKHVNNF
                :*  * :  *. .**::*...: ** :::**:  :   : ::.:***   *. ::: .       ::  *:.::   : :*

AAM76037.1      INKLDALHVVNYNGLLSSIEIGTQNHTIIITRTNMGFLVELQEPDKSAM--NRMKPGPAKFSLLHESTLKAFTQGSSTRM
ACI28626.1      ISQLDALHVVNYNGLLSSIEIGTRNHTIIITRTNMGFLVELQEPDKSAM--NQKKPGPVKFSLLHESTFKALIKKPATKM
ACI28635.1      INQLDALHVVNYNGLLSSIEIGTKSHTIIITRTNMGFLVELQEPDKSAM--NTRKPGPVKFSLLHESTLKTLAKKPATQM
AGL50930.1      INKLETLHVVNYKGLLSSVEIGTPSYAIIITRTNMGYLVEVQEPDKSAM--DIRHPGPVKFSLLHESTLKPVATPKPSSI
CAA78119.1      FNKMFTLHVVNDHGKPSSIEIKLTGQQIIITRVNMGFLVEVRRIDIEPCCGETVLSESVVFGLVAEAVLREHSQMEKGQP
UKR35333.1      ILKLLELHVINNRGFTSSIEIGNKYNTIIISRTNMGYLVEQQEPDKSAL--RKTKPGPVKFTLVHEDAFRDFK-PERSSI
                : ::  ***:* .*  **:**      ***:*.***:*** :. * ..       . .. * *: * .::

AAM76037.1      QSLILEFNSSLAI
ACI28626.1      QALILEFNSSLAI
ACI28635.1      QALILEFNSSLAI
AGL50930.1      TSLIMEFNSSLAI
CAA78119.1      LNLTQYMNSKIAI
UKR35333.1      NLLIMEFNSSLAI
                  *   :**.:**
```

AAM76037.1 VP24 [Zaire ebolavirus]
ACI28626.1 VP24 [Bundibugyo ebolavirus]
ACI28635.1 VP24 [Tai Forest ebolavirus]
AGL50930.1 VP24 [Sudan ebolavirus]
CAA78119.1 vp24 [Marburg virus - Musoke, Kenya, 1980]
UKR35333.1 VP24 [Cuevavirus lloviuense]

## C VP35

```
AAM76032.1      MTT-RTKGRGHTVATTQNDRMPGPELSGWISEQLMTGRIPVSDIFCD--IENNPG--------LCYASQM-QQTKPNPKT
ACI28621.1      MTSNRARVTYNPPPTTTGTRSCGPELSGWISEQLMTGKIPITDIFNE--IETLPS--------ISPSIHS-KIKTPSVQT
ACI28630.1      MISTRAAAINDPSLPIRNQCTRGPELSGWISEQLMTGKIPVHEIFND--TEPHIS--------SGSDCLP-RPKNTAPRT
ACR33188.1      MQQDKT------------YRHHGPEVSGWFSEQLMTGKIPLTEVFVD--VETKPS--------PTPITII-SKNPKTTRK
CAA78115.1      MWD--------------------SSYMQQVSEGLMTGKVPIDQVFGANPLEKLYK-RRKPKGTVGL--------QCSPCL
UKR35327.1      MD-------------------------KVTDAIMTGRLSLDTIPGF--QCSAPTSRSRP---TGRKCPPWEGPLIEPKT
                *                        .:: :***::.:  :

AAM76032.1      -RNSQTQTDPICNHSFEEVVQTLASLATVVQQQTIASES----LEQRITSLEN---GLKPVYDMAKTISSLNRVCAEMVA
ACI28621.1      -RSVQTQTDPNCNHDFAEVVKMLTSLTLVVQKQTLATES----LEQRITDLEG---SLKPVSEITKIVSALNRSCAEMVA
ACI28630.1      -RNTQTQTDPVCNHNFEDVTQALTSLTNVIQKQALNLES----LEQRIIDLEN---GLKPMYDMAKVISALNRSCAEMVA
ACR33188.1      -SDKQVQTDDASSLLTEEVKTAINSVISAVRRQTNAIES----LESRIANLEA---SLKPVQDMAKTISSLNRSCAEMVA
CAA78115.1      -MSKATSTDDIIWD-QLIVKRTLADLLIPINRQISDIQSTLSEVTTRVHEIERQLHEITPVLKMGRTLEAISKGMSEMLA
UKR35327.1      SANKSTQTENIYQ--SDQVLREIKQLSSILAHQITTMER----LEGSLQSIHQ---QLQVVGGMAHNLTSLTTLCTEMVA
                  .  ..*:         *   : .:   : :*    :     :   : .:.      :  :  : : : ::.   :**:*

AAM76032.1      KYDLLVMTTGRATATAAATEAYWAEHGQPPPGPSLYEESAIRGKIESRDETVPQSVREAFNNLNSTTSLTEENFGKPDIS
ACI28621.1      KYDLLVMTTGRATATAAATEAYWAEHGRPPPGPSLYEEDAIRTKIGKQGDMVPKEVQEAFRNLDSTALLTEENFGKPDIS
ACI28630.1      KYDLLVMTTGRATATAAATEAYWEEHGQPPPGPSLYEESAIRGKINKQEDKVPKEVQEAFRNLDSTSSLTEENFGKPDIS
ACR33188.1      KYDLLVMTTGRATATAAATEAYWNEHGQAPPGPSLYEDDAIKAKLKDPNGKVPESVKQAYTNLDSTSALNEENFGRPYIS
CAA78115.1      KYDHLVISTGRTTAPAAAFDAYLNEHGVPPPQPAIFKDLGVAQQACSKGTMVKNATTDAADKMSKVLELSEETFSKPNLS
UKR35327.1      KYDFLVMTTGRATATAAATEAYWKEHGKAPPGPALFEADALKTRIQENPKSVPTDVREAFDRLEKTEEVTERTFGKPTIS
                *** **::***:**.*** :**   *** .** *::::  .:  :  .     *   . :*  .:...  :.*..*.:* :*

AAM76032.1      AKDLRNIMYDHLPGFGTAFHQLVQVICKLGKDSNSLDIIHAEFQASLAEGDSPQCALIQITKRVPIFQDAAPPVIHIRSR
ACI28621.1      AKDLRNIMYDHLPGFGTAFHQLVQVICKLGKDNSSLDVIHAEFQASLAEGDSPQCALIQITKRIPIFQDAAPPVIHIRSR
ACI28630.1      AKDLRDIMYDHLPGFGTAFHQLVQVICKLGKDNSALDIIHAEFQASLAEGDSPQCALIQITKRIPIFQDATPPTIHIRSR
ACR33188.1      AKDLKEIIYDHLPGFGTAFHQLVQVICKIGKDNNILDIIHAEFQASLAEGDSPQCALIQITKRIPTFQDASPPIVHIKSR
CAA78115.1      AKDLALLLFTHLPGNNTPFHILAQVLSKIAYKSGKSGAFLDAFHQILSEGENAQAALTRLSRTFDAFLGVVPPVIRVKNF
UKR35327.1      AKLLKELFYDHLPGYGTAFHQLAQVVCKVGKDNDLLDIIHAEFQASLAEGDSPQCALIQITKRIPIFGETPPPVIYIKTK
                ** *  ::: ****  .*.** *.**:.*:. ...   . :    *:  *:**:..*.** :::: .   *  . ** : ::.

AAM76032.1      GDIPRACQKSLRPVPPSPKIDRGWVCVFQLQDGKTLGLKI
ACI28621.1      GDIPKACQKSLRPVPPSPKIDRGWVCIFQLQDGKTLGLKI
ACI28630.1      GDIPRACQKSLRPVPPSPKIDRGWVCIFQLQDGKTLGLKI
ACR33188.1      GDIPKACQKSLRPVPPSPKIDRGWVCIFQFQDGKTLGLKI
CAA78115.1      QTVPRPSQKSLRAVPPNPTIDKGWVCVYSSEQGETRALKI
UKR35327.1      QEIPKACQKSLRPLPPNPKIERGWVCHYVTADGKEMGLKI
                  :*:..*****.:**.*.*::**** :   :*:  .***
```

AAM76032.1 VP35 [Zaire ebolavirus]
ACI28621.1 VP35 [Bundibugyo ebolavirus]
ACI28630.1 VP35 [Tai Forest ebolavirus]
ACR33188.1 VP35 [Sudan ebolavirus]
CAA78115.1 VP35 [Marburg virus - Musoke, Kenya, 1980]
UKR35327.1 VP35 [Cuevavirus lloviuense]

**Figure S6 (continue). Sequence alignment of orthologs of NP, VP24, and VP35 and regions involved in the interface and average conservation scores**

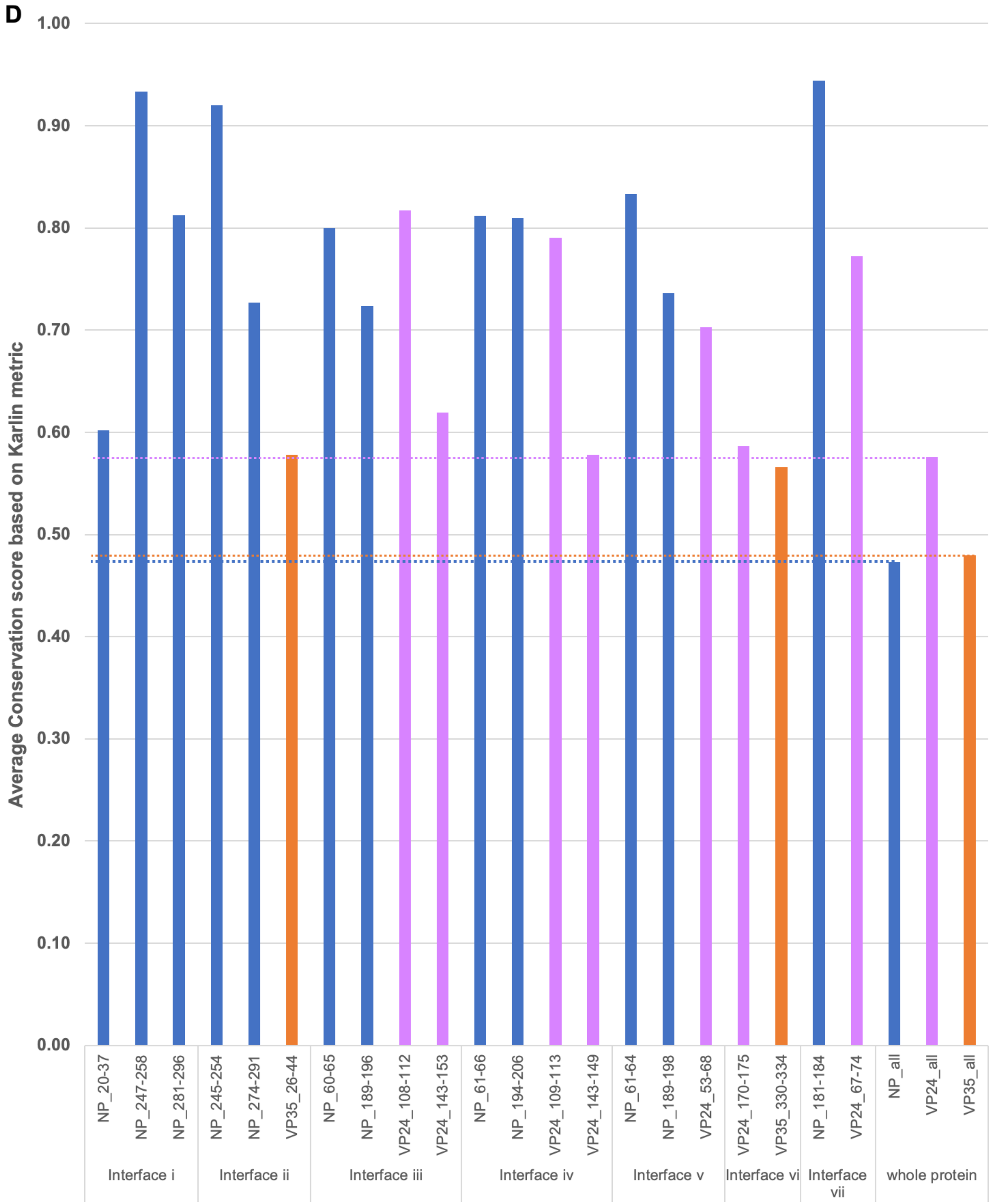

**Figure S7. Potential location of VP35 oligomerization domain**

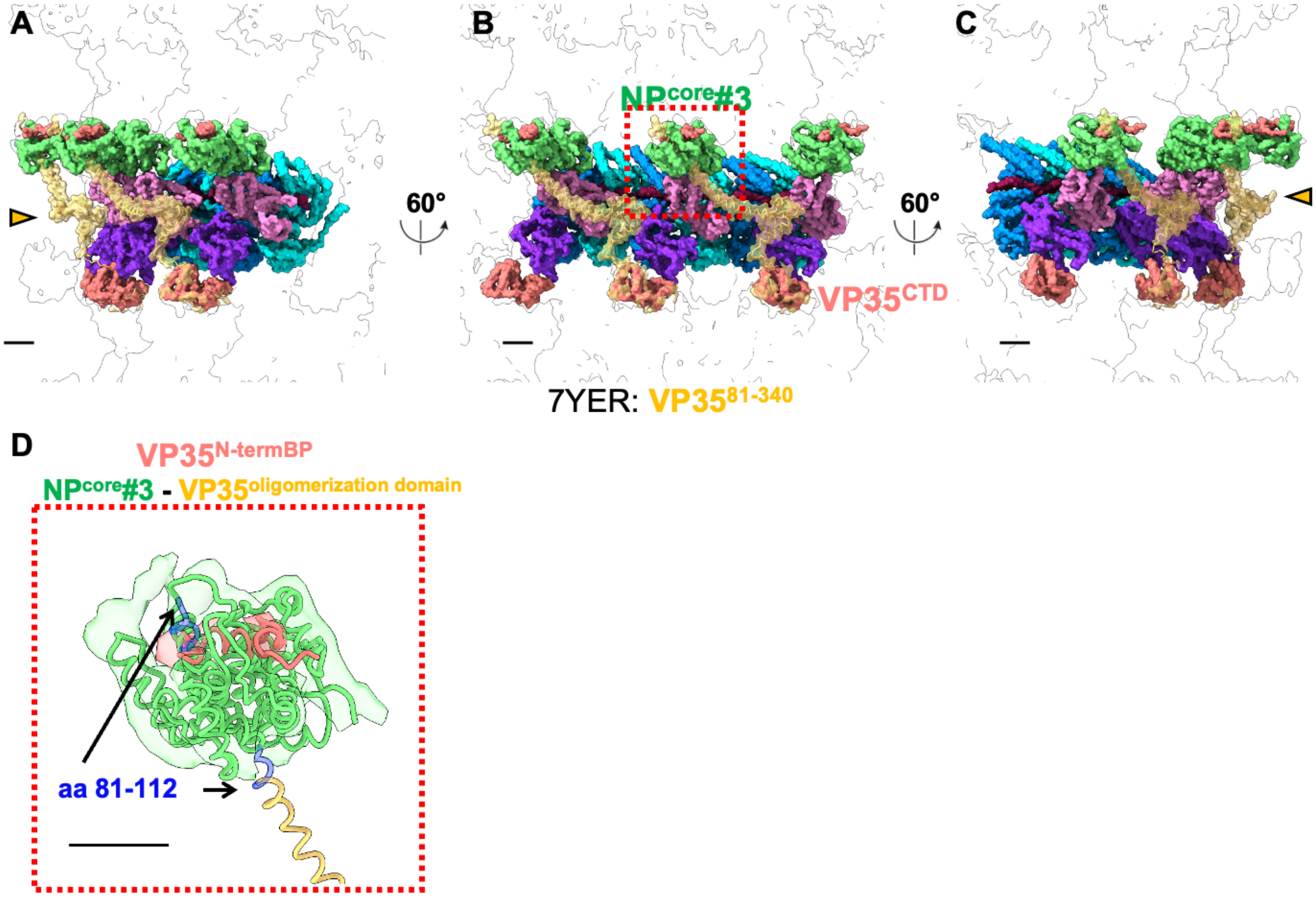

**Table S1. Interaction sites involved in nucleocapsid assembly and average conservation scores obtained with three different scoring metrics, related to Figure 6**

| | Protein_regions (amino acid) | Karlin conservation score | Livingstone conservation score | Valdar conservation score |
|---|---|---|---|---|
| Interface (**i**) NP#1-NP#2 | NP_20-37 | 0.60 | 7.56 | 7.02 |
| | NP_247-258 | 0.93 | 10.50 | 8.20 |
| | NP_281-296 | 0.81 | 9.19 | 7.61 |
| Interface (**ii**) $VP35^{N\text{-}termBP}$-NP#3 | NP_245-254 | 0.92 | 10.40 | 8.17 |
| | NP_274-291 | 0.73 | 8.56 | 7.29 |
| | VP35_26-44 | 0.58 | 7.21 | 6.93 |
| Interface (**iii**) NP#1-VP24#1 | NP_60-65 | 0.80 | 9.00 | 7.64 |
| | NP_189-196 | 0.72 | 8.50 | 7.32 |
| | VP24_108-112 | 0.82 | 8.80 | 12.09 |
| | VP24_143-153 | 0.62 | 7.64 | 11.19 |
| Interface (**iv**) VP24#2-NP#3 | NP_61-66 | 0.81 | 9.00 | 7.76 |
| | NP_194-206 | 0.81 | 9.15 | 7.73 |
| | VP24_109-113 | 0.79 | 8.00 | 11.73 |
| | VP24_143-149 | 0.58 | 7.43 | 11.03 |
| Interface (**v**) NP#2-VP24#2 | NP_61-64 | 0.83 | 8.75 | 7.81 |
| | NP_189-198 | 0.74 | 8.60 | 7.38 |
| | VP24_53-68 | 0.70 | 8.25 | 11.40 |
| Interface (**vi**) VP24#1-$VP35^{CTD}$ | VP24_170-175 | 0.59 | 7.67 | 10.83 |
| | VP35_330-334 | 0.57 | 6.60 | 7.18 |
| Interface (**vii**) VP24#1-inter-rung NP#3 | NP_181-184 | 0.94 | 10.75 | 8.17 |
| | VP24_67-74 | 0.77 | 8.75 | 11.68 |
| whole protein | NP_all | 0.47 | 6.44 | 6.11 |
| | VP24_all | 0.58 | 7.54 | 10.59 |
| | VP35_all | 0.48 | 6.48 | 6.26 |

**Table S2. Cryo-ET data collection and processing, related to Figures 1, 2, 3, and 5**

| | Intracellular Ebola NPΔ601-739-VP24-VP35 nucleocapsid | Intracellular Ebola NP-VP24-VP35 nucleocapsid | Intracellular EBOV-GFP-ΔVP30 nucleocapsid | EBOV-GFP-ΔVP30 nucleocapsid |
|---|---|---|---|---|
| **Data collection and processing** | | | | |
| Microscope | Krios | | | |
| Voltage | 300 kV | | | |
| Detector | Gatan K3 | | | |
| Energy filter | BioQuantum energy filter | | | |
| Slit width (eV) | 20 | | | |
| Defocus (μm) | 2.5 to 3.5 | 4.5 to 5.5 | | 2.5 to 4.5 |
| Pixel size (Å) | 2.1488 | 2.6848 | | 1.7 |
| Number of frames | 10 | | | |
| Numbers of Tilts | 41 | | | |
| Tilt range | -40 to +40 in 2˚increments | -60 to +60 in 3˚increments | | |
| Acquisition Scheme | Dose-symmetric | | | |
| Electron exposure ($e^-$/Å$^2$) | 140-160 | | | |
| Number of tomograms used | 25 | 10 | - | |
| Symmetry imposed | Not applied | | - | |
| Total length of nucleocapsid (μm) | 220 | 128 | - | |
| Final no. of particles | 28'273 | 8'371 | - | |
| Helical rise (Å) | 7.35 | 7.18 | - | |
| Helical twist (°) | -30.35 | -30.45 | - | |
| Map resolution at 0.143 FSC (Å) | 9.37 | 17.57 | - | |
| **EMDB** | **EMD-42509** | **EMD-42515** | - | |

**Table S3. Refinement statistics, related to Figures 2, and 4**

| | Intracellular Ebola virus nucleocapsid | In virion Ebola virus nucleocapsid |
|---|---|---|
| Refinement | | |
| **R.M.S. Deviations** | | |
| Bond Length (Å) | 0.004 | 0.004 |
| Bond Angles(˚) | 1.102 | 1.162 |
| **Validation** | | |
| Molprobity score | 1.66 | 1.65 |
| Poor rotamers (%) | 0 | 0 |
| **Ramachandran plot** | | |
| Outliers (%) | 0 | 0.26 |
| Allowed (%) | 7.76 | 6.95 |
| Favored (%) | 92.24 | 92.79 |
| **PDB** | **8USN** | **8UST** |
